# Cholinergic activation of corticofugal circuits in the adult mouse prefrontal cortex

**DOI:** 10.1101/2023.04.28.538437

**Authors:** Allan T. Gulledge

**Affiliations:** Department of Molecular and Systems Biology, Geisel School of Medicine at Dartmouth College 74 College Street, Vail 601, Hanover, New Hampshire 03755, USA

**Keywords:** Prefrontal cortex, prelimbic cortex, acetylcholine, muscarinic receptor, pyramidal neuron, mouse

## Abstract

In layer 5 of the neocortex, ACh promotes cortical output to the thalamus and brainstem by preferentially enhancing the postsynaptic excitability of pyramidal tract (PT) neurons relative to neighboring intratelencephalic (IT) neurons. Less is known about how ACh regulates the excitatory synaptic drive of IT and PT neurons. To address this question, spontaneous excitatory postsynaptic potentials (sEPSPs) were recorded in pairs of IT and PT neurons in slices of prelimbic cortex from adult female and male mice. ACh (20 µM) enhanced sEPSP amplitudes, frequencies, rise-times, and half-widths preferentially in PT neurons. These effects were blocked by the muscarinic acetylcholine receptor antagonist atropine (1 µM). When challenged with pirenzepine (1 µM), an antagonist selective for M1-type muscarinic receptors, ACh instead reduced sEPSP frequencies. The cholinergic increase in sEPSP amplitudes and frequencies in PT neurons was not sensitive to blockade of GABAergic receptors with gabazine (10 µM) and CGP52432 (2.5 µM), but was blocked by tetrodotoxin (1 µM), suggesting that ACh enhances action-potential-dependent excitatory synaptic transmission in PT neurons. ACh also preferentially promoted the occurrence of synchronous sEPSPs in pairs of PT neurons relative to IT-PT and IT-IT pairs. Finally, selective chemogenetic silencing of hM4Di-expressing PT, but not IT, neurons with clozapine-N-oxide (5 µM) blocked cholinergic enhancement of sEPSP amplitudes and frequencies in PT neurons. These data suggest that, in addition to enhancing the postsynaptic excitability of PT neurons, M1 receptor activation promotes corticofugal output by preferentially amplifying recurrent excitation within networks of PT neurons.

## Introduction

Output from the neocortex is segregated into two broad, non-overlapping channels based on the axonal projection patterns of pyramidal neuron subpopulations: pyramidal tract (PT) neurons provide corticofugal output to deep subcortical structures, such as the thalamus and brainstem, whereas most corticocortical projections arise from intratelencephalic (IT) projecting neurons (for review, see Baker et al., 2018a). These pyramidal neuron subpopulations exist side-by-side in layer 5 of the neocortex, including the prelimbic cortex, an area of medial prefrontal cortex (mPFC) critical for decision making and goal-directed behavior (for reviews, see Sharpe et al., 2019; Green and Bouton, 2021).

A growing literature has revealed that IT and PT neurons are differentially sensitive to neuromodulators allowing for brain-state-dependent regulation of cortical circuit output (e.g., Dembrow et al., 2010; e.g., Avesar and Gulledge, 2012; Gee et al., 2012; Elliott et al., 2018). Experiments in this laboratory and others have found that ACh preferentially enhances the intrinsic excitability of PT neurons, relative to IT neurons, in juvenile (Joshi et al., 2016) and adult rodents (Dembrow et al., 2010; Baker et al., 2018b) via activation of postsynaptic M1-type muscarinic acetylcholine receptors (mAChRs; Gulledge et al., 2009). Yet, rather than directly exciting PT neurons, ACh increases postsynaptic gain to amplify action potential output in response to excitatory synaptic drive (Andrade, 1991; Gulledge and Stuart, 2005; Williams and Fletcher, 2019). Therefore, understanding the net effect of ACh on cortical circuit output will require determination of how ACh regulates excitatory synaptic transmission in IT and PT neurons.

Prior studies have found that ACh acts presynaptically to suppress glutamate release at many synapses in the neocortex (Vidal and Changeux, 1993; Gil et al., 1997; Tsodyks and Markram, 1997; Hsieh et al., 2000; Atzori et al., 2005; Levy et al., 2006) and hippocampus (Hounsgaard, 1978; Valentino and Dingledine, 1981; Segal, 1982; Hasselmo and Schnell, 1994; Hasselmo et al., 1995; Fernandez de Sevilla et al., 2002; Goswamee and McQuiston, 2019). The inhibitory effect of ACh on excitatory synaptic transmission is attributable primarily to activation of M4-type mAChRs (Kimura and Baughman, 1997; Shirey et al., 2008; Eggermann and Feldmeyer, 2009; Dasari and Gulledge, 2011; Yang et al., 2020). On the other hand, ACh may enhance glutamate release at cortical synapses via presynaptic nicotinic acetylcholine receptors (nAChRs; Gil et al., 1997; Aramakis and Metherate, 1998; Urban-Ciecko et al., 2018; Yang et al., 2020). ACh may also increase (Yang et al., 2020) or decrease (Atzori et al., 2005) the frequency of spontaneous “miniature” glutamatergic synaptic events in neocortical neurons, and can increase the frequency and amplitude of action-potential-dependent spontaneous excitatory events in layer 5 neurons of the mPFC (Shirey et al., 2009).

Aside from the substantial evidence for muscarinic suppression of evoked transmitter release at cortical synapses, results in the studies described above are variable and inconsistent. This may reflect developmental differences in cholinergic signaling (e.g., Aramakis and Metherate, 1998), as most of the studies described above utilized neurons spanning from young to adolescent animals (8-days old to ∼4 weeks old). Alternatively, variation may arise from differences in cortical areas studied, species differences (e.g., mice vs rats; see Elliott et al., 2018), or differences in other experimental parameters. Of note, only one prior study tested for projection-specific cholinergic modulation of excitatory drive (Yang et al., 2020), doing so in two populations of layer 6 projection neurons from immature (≤ 3 weeks old) animals.

This study was designed to compare the impact of ACh on the net excitatory drive of layer 5 IT and PT neurons in the adult mPFC. Given the considerable evidence for muscarinic suppression of glutamate release across diverse experimental conditions, it was hypothesized that ACh should reduce both the frequency and amplitudes of spontaneous excitatory postsynaptic potentials (sEPSPs) in IT and PT neurons. As detailed below, this hypothesis was tested using simultaneous recordings of sEPSPs in pairs of IT and PT neurons in slices of the adult mouse prelimbic cortex.

## Methods

### Animals and ethical approvals

Experiments were performed using female and male 6- to 16-week-old (mean ± SD of 10.4 ± 2.5 weeks-old, n = 51 mice) C57BL/6J wild-type mice cared for under protocols approved by the Institutional Animal Care and Use Committee of Dartmouth College. Animals were bred and maintained on a 12h light-dark cycle with free access to food and water in facilities accredited by the Association for Assessment and Accreditation of Laboratory Animal Care.

### Animal surgeries

For some experiments, an AAV retrograde virus expressing pAAV-hSyn-hM4D(Gi)-mCherry (Addgene #50475) or pAAV-hSyn-hM3D(Gq)-mCherry (Addgene #50474) was injected into either the ipsilateral pons (relative to lambda, 1.00 mm lateral, 0.20 mm posterior, depth of 4.55 mm) or contralateral mPFC (relative to bregma, 0.48 mm lateral, 2.1 mm anterior, depth of 1.6 mm) of 6- to 9-week-old mice to express DREADD (“designer receptor exclusively activated by designer drugs”) receptors selectively in PT (ipsilateral pons injections) or IT (contralateral mPFC injections) neurons in the prefrontal cortex. Under sterile surgical conditions, animals were anesthetized with continuous isoflurane (∼2%) and a craniotomy made at the above coordinates. A 33 gauge microsyringe (Hamilton) containing undiluted virus was slowly lowered into place over a period of ∼5 minutes. After waiting anther 5 minutes, the virus was injected at a rate of 50 nL per minute for a total volume of 450 nL (pons) or 300 nL (mPFC). Five minutes after the injection was completed, the microsyringe was slowly removed and the wound sutured. Mice were allowed to recover from surgery for at least 21 days before use in experiments.

### Slice preparation

Mice were anesthetized with vaporized isoflurane and decapitated. Brains were rapidly removed and sliced in an artificial cerebral spinal fluid (aCSF) composed of (in mM): 125 NaCl, 25 NaHCO_3_, 3 KCl, 1.25 NaH_2_PO_4_, 0.5 CaCl_2_, 6 MgCl_2_, 0.01 ascorbic acid, and 25 glucose (saturated with 95% O_2_ / 5% CO_2_). Coronal brain slices (250 µm thickness) containing the mPFC were stored in a holding chamber filled with normal recording aCSF containing 2 mM CaCl_2_ and 1 mM MgCl_2_ for 1 hour at 35 °C, and then maintained at room temperature (∼26 °C) until use in experiments.

### Electrophysiology

Slices were transferred to a recording chamber (∼0.5 ml volume) perfused continuously (∼6 ml/min) with oxygenated aCSF heated to 35-36 °C. Layer 5 pyramidal neurons in the prelimbic cortex were visualized using oblique illumination with a 60x water-immersion objective (Olympus). Electrical recordings were made of layer 5 neurons using patch pipettes (5-7 MΩ) filled with a solution containing (in mM): 140 K-gluconate, 2 NaCl, 2 MgCl_2_, 10 HEPES, 3 Na_2_ATP, and 0.3 NaGTP, pH 7.2 with KOH. Data were acquired in current-clamp using a 2-electrode amplifier (dPatch, Sutter Instrument) connected to a Mac Studio computer (Apple, Inc.) running SutterPatch software under Igor Pro (Sutter Instrument). Capacitance was maximally neutralized and series resistances (typically 15-25 MΩ) were compensated with bridge balance. Membrane potentials were sampled at 25 kHz (250 kHz for action potential waveforms), filtered at 5 kHz (50 kHz for action potentials), and corrected for the liquid junction potential (+12 mV).

Neurons were classified as either IT or PT based on their physiological profiles (**Table 1**, **Figure 1**), as quantified by a Physiology Index (PI) that successfully identified 93% of retrograde-labeled projection neurons from an earlier study (Baker et al., 2018b; **see Figure 1D**). Similar physiological measures have been used by others to sort IT and PT neurons in the mPFC (Lee et al., 2014; Elliott et al., 2018; Meda et al., 2019). The PI was computed for each neuron based on the magnitude of the HCN-channel-mediated rebound “sag” potential (as a percent of the peak hyperpolarization following a current pulse sufficient to generate a peak hyperpolarization of ∼20 mV below the resting membrane potential), the “slope of sag” rebound (as measured by linear regression over a 50 ms period beginning at the time of peak hyperpolarization), and the input resistance (R_N_) of the neuron, according to the following formula:

**Figure 1.**
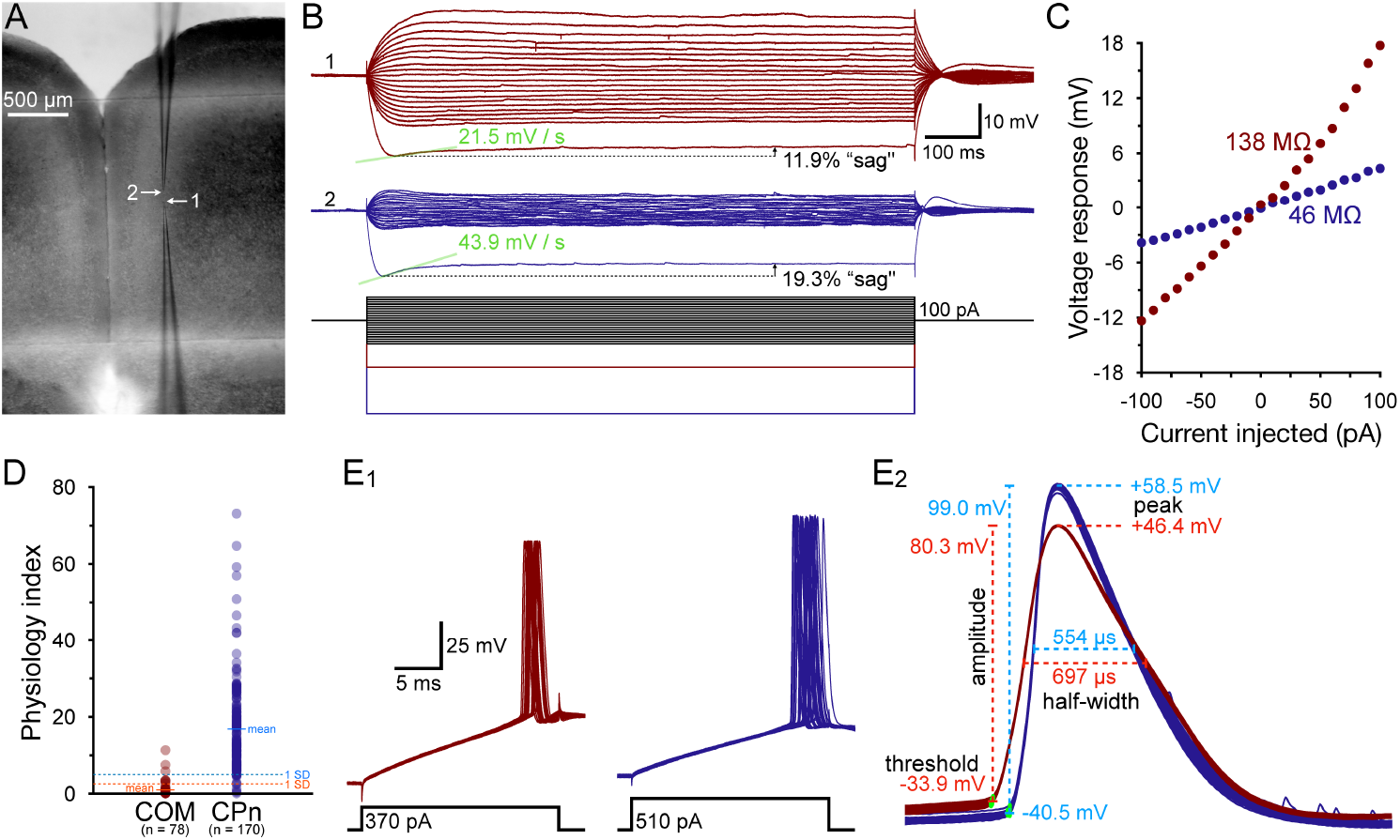
Classification of intratelencephalic (IT) and pyramidal tract (PT) layer 5 projection neurons. ***A***) Image of a coronal slice of mouse brain containing the medial prefrontal cortex (5x objective) showing the locations of recording electrodes in layer 5. ***B***) Voltage responses in the two neurons recorded in ***A*** (top and middle traces) to a series of current steps (lower traces). Note that neuron 1 (red) has **a**) larger voltage deflections to current steps (bottom traces) and a correspondingly larger input resistance, **b**) a smaller depolarizing “sag” response after a 20 mV hyperpolarizing step (indicative of limited HCN channel activation), and **c**) a smaller slope of the initial sag response than observed in neuron 2 (blue). These values were used to compute a Physiology Index (PI; see Methods) of 1.85 and 18.6 for neurons 1 and 2, respectively. ***C***) Plots of the voltage-current relationships for data shown in ***B***. Linear regressions from -50 to +50 pA were used to calculate the input resistances of these neurons. ***D***) PIs calculated for 248 retrograde-labeled commissural projection (COM) neurons (a type of IT neuron) and corticopontine (CPn) neurons (a type of PT neuron) from Baker et. al. (2018). Labeled COM neurons (n = 78) had a mean (± SD) PI of 0.85 ± 1.72, whereas labeled CPn neurons (n = 170) had a mean PI of 16.9 ± 11.9. In this study, cutoffs of 1 standard deviation from the mean (2.6 or 5.0) were used to physiologically classify IT and PT neurons, respectively, with neurons with PIs between 2.6 and 5 remaining unclassified and excluded from analysis. This approach correctly classified 92.7% of neurons from Baker et al., and misclassified only 3.6% of neurons, with the remainder being left unclassified (PIs of between 2.6 and 5.0). ***E_1_***) Brief (20 ms) current steps were used to evoke action potentials captured at 250 kHz (30 superimposed trials) in the IT and PT neurons shown in ***B*** and ***C***. ***E_2_***) Magnification of the peak-aligned action potentials shown in ***E_1_***, showing measurements made of spike thresholds, amplitudes, peaks, and widths at half-amplitude, as listed in **Table 2**.

**Table 1.** Physiological classification of IT and PT neurons for each experiment.

| Neuron group | Neuron type | n | R<sub>N</sub> (MΩ) | Sag (%) | Sag slope (mV/s) | Physiology index |
| --- | --- | --- | --- | --- | --- | --- |
| **Figure 3** | IT | 19 | 165 ± 84 | 7.6 ± 3.6 | 13 ± 8 | 0.7 ± 0.6 |
|  | PT | 19 | 56 ± 12 | 18.7 ± 4.5 | 50 ± 16 | 18.5 ± 11.7 |
| **Figure 4** | PT | 10 | 58 ± 15 | 19.2 ± 3.8 | 50 ± 15 | 18.6 ± 10.2 |
| **Figure 5** | PT | 13 | 49 ± 12 | 16.0 ± 3.0 | 38 ± 10 | 13.3 ± 5.8 |
| **Figure 6** | PT | 13 | 45 ± 11 | 16.9 ± 4.6 | 38 ± 10 | 15.9 ± 9.8 |
| **Figure 7** | PT | 16 | 67 ± 17 | 21.3 ± 6.2 | 49 ± 16 | 17.6 ± 12.1 |
| **Figure 8** | IT | 24 | 145 ± 57 | 9.2 ± 3.5 | 16 ± 9 | 1.2 ± 0.9 |
|  | PT | 44 | 57 ± 15 | 17.6 ± 4.1 | 46 ± 16 | 15.6 ± 8.5 |
| **Figure 9** | PT* | 21 | 51 ± 15 | 18.1 ± 4.8 | 42 ± 15 | 17.0 ± 10.8 |
| **Figure 10** | PT | 12 | 53 ± 7 | 17.3 ± 3.3 | 36 ± 5 | 12.1 ± 3.8 |
| **Figure 11** | PT* | 9 | 43 ± 11 | 16.9 ± 3.9 | 34 ± 11 | 14.5 ± 6.4 |
| **All neurons** | IT | 43 | 154 ± 70 | 8.5 ± 3.6 | 15 ± 9 | 1.0 ± 0.8 |
|  | PT | 157 | 54 ± 15 | 18.0 ± 4.5 | 44 ± 15 | 16.0 ± 9.4 |

Data are shown as means ± SD. \*Targeted PT neurons retrograde-labeled with hM4Di- or hM3Dq-mCherry. Statistical treatments are not applicable, as IT and PT populations were defined based on measurements of R<sub>N</sub>, % sag, and sag slope (i.e., the Physiology Index; see **Methods**)

**Table 2.** Additional physiological properties of IT and PT neurons in each experiment.

| Neuron group | Neuron type | n | Resting $V_m$ (mV) | AHP amplitude (mV) | AP threshold (mV) | AP peak (mV) | AP amplitude (mV) | AP half-width (ms) |
| --- | --- | --- | --- | --- | --- | --- | --- | --- |
| Figure 3 | IT | 19 | $-83.6 \pm 3.8$ | $-6.3 \pm 3.1$ | $-49.3 \pm 4.3$ | $+44.5 \pm 6.1$ | $93.8 \pm 7.5$ | $604 \pm 104$ |
| | PT | 19 | $-79.5 \pm 2.7$ | $-7.1 \pm 1.2$ | $-54.1 \pm 3.7$ | $+45.7 \pm 8.0$ | $103.8 \pm 6.7$ | $558 \pm 61$ |
|  | IT vs PT t-test<sup>†</sup> (p) |  | < 0.001 | 0.269 | < 0.001 | 0.088 | 0.001 | 0.068 |
|  | Effect size (d) |  | **1.26** | 0.35 | **1.19** | **0.79** | **1.41** | 0.53 |
| Figure 4 | PT | 10 | $-80.4 \pm 2.9$ | $-7.1 \pm 0.7$ | $-55.1 \pm 3.0$ | $+48.5 \pm 3.9$ | $103.6 \pm 4.8$ | $524 \pm 32$ |
| Figure 5 | PT | 13 | $-80.1 \pm 2.4$ | $-6.6 \pm 1.2$ | $-53.2 \pm 2.6$ | $+46.5 \pm 4.6$ | $98.4 \pm 6.4$ | $507 \pm 70$ |
| Figure 6 | PT | 13 | $-79.1 \pm 2.5$ | $-7.1 \pm 1.1$ | $-55.5 \pm 2.2$ | $+46.4 \pm 2.6$ | $101.8 \pm 3.3$ | $460 \pm 53$ |
| Figure 7 | PT | 16 | $-77.7 \pm 2.4$ | $-7.1 \pm 1.1$ | $-52.4 \pm 3.7$ | $+45.9 \pm 7.6$ | $98.3 \pm 8.6$ | $565 \pm 70$ |
| Figure 8 | IT | 24 | $-82.5 \pm 3.2$ | $-5.8 \pm 1.4$ | $-47.0 \pm 4.9$ | $+43.9 \pm 6.4$ | $90.9 \pm 6.8$ | $686 \pm 112$ |
| | PT | 44 | $-78.9 \pm 3.2$ | $-6.6 \pm 1.6$ | $-52.4 \pm 3.2$ | $+47.0 \pm 6.3$ | $99.3 \pm 7.3$ | $553 \pm 79$ |
|  | IT vs PT t-test<sup>†</sup> (p) |  | < 0.001 | 0.033 | < 0.001 | 0.065 | < 0.001 | < 0.001 |
|  | Effect size (d) |  | **1.30** | 0.60 | **1.54** | 0.55 | **1.33** | **1.64** |
| Figure 9 | PT* | 21 | $-79.2 \pm 2.3$ | $-6.9 \pm 0.7$ | $-54.9 \pm 2.0$ | $+44.3 \pm 4.2$ | $99.1 \pm 4.4$ | $459 \pm 59$ |
| Figure 10 | PT | 12 | $-81.4 \pm 3.3$ | $-7.7 \pm 1.3$ | $-55.8 \pm 2.9$ | $+43.2 \pm 3.8$ | $99.0 \pm 4.8$ | $478 \pm 44$ |
| Figure 11 | PT* | 9 | $-76.9 \pm 4.8$ | $-7.5 \pm 1.0$ | $-55.2 \pm 1.6$ | $+44.7 \pm 2.7$ | $100.0 \pm 2.8$ | $435 \pm 32$ |
| All neurons | IT | 43 | $-83.0 \pm 3.5$ | $-6.0 \pm 2.3$ | $-48.0 \pm 4.7$ | $+44.1 \pm 6.2$ | $92.1 \pm 7.2$ | $652 \pm 115$ |
| | PT | 157 | $-79.9 \pm 3.8$ | $-7.0 \pm 1.3$ | $-53.8 \pm 3.2$ | $+46.3 \pm 5.6$ | $100.1 \pm 6.3$ | $515 \pm 77$ |
|  | IT vs PT t-test<sup>†</sup> (p) |  | < 0.001 | 0.013 | < 0.001 | 0.046 | < 0.001 | < 0.001 |
|  | Effect size (d) |  | **0.87** | 0.61 | **1.65** | 0.38 | **1.22** | **1.58** |

Data presented as means ± SD. \*Targeted PT neurons retrograde-labeled with hM4Di- or hM3Dq-mCherry. <sup>†</sup>Student's t-tests for paired samples were used for comparisons of means of physiological attributes for paired recordings from IT and PT neurons. <sup>†</sup>Student's t-tests assuming unequal variances were used for comparisons of physiological attributes of IT and PT neurons recorded as homotypic pairs and for the combined data sets. Large effect sizes ( $\geq \sim 0.80$ ) are shown in bold.

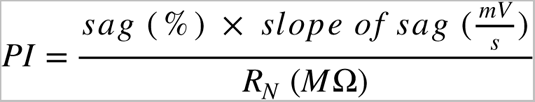

When calculated for retrograde-labeled neurons studied in a previous paper (Baker et al., 2018b), layer 5 IT neurons projecting to the contralateral hemisphere (n = 78) had a mean (± SD) PI of 0.85 ± 1.72, while labeled pons-projecting PT neurons (n = 170) had a mean PI of 16.9 ± 11.9. In this study, neurons were classified as “IT” if their calculated PI was less than 2.6 (one SD above the mean for labeled IT neurons in Baker et al., 2018b) or “PT” if their PIs were greater than 5.0 (one SD below the mean of labeled PT neurons in the previous study). Neurons with PIs between 2.6 and 5.0 were considered ambiguous and excluded from analysis. When tested against labeled commissural (IT) and brainstem projecting (PT) neurons from another previous paper (Avesar and Gulledge, 2012), this classification scheme misidentified only 1 of 43 (2.3%) neurons. The classification scheme also correctly identified all 30 retrograde-labeled PT neurons studied in this present report (**Table 1**).

For each neuron, additional physiological measurements were made of the resting membrane potential, the amplitude of afterhyperpolarizations (AHPs), and action potential (AP) attributes, including AP threshold (as detected with a 50 mV/ms threshold), peak voltages, amplitudes from threshold, and widths at half-ampli-tudes (i.e., half-widths; **Table 2**, **Figure 1E**). AHPs were generated with a train of brief (2-ms) high amplitude (3 nA) current pulses delivered at 50 Hz for three seconds (Gulledge et al., 2013), and peak AHPs were measured relative to initial resting membrane potentials. APs were evoked with 20-ms current pulses typically in the range of 300 - 500 pA.

Following baseline physiological measurements, spontaneous excitatory postsynaptic potentials (sEPSPs) were recorded over 30 minutes with neurons “held” at their resting membrane potentials using the dPatch “dynamic holding” function that introduces a slow bias current to counteract changes in resting membrane potential due to ACh or other drugs (e.g., clozapine-n-oxide) even as much faster synaptic potentials are faithfully recorded. Quantifying the ACh-induced bias current over time also provided a measure of the postsynaptic effects of ACh or other substances on IT and PT neurons. During recordings, ACh and/or other substances were bath-applied for periods of time (typically 7 minutes for bath-applied ACh). Data were analyzed offline using Axograph or Igor Pro. Each 30-minute sweep of spontaneous activity was high-pass filtered at 0.2 Hz, low-pass filtered at 5 kHz, and notch-filtered at 60 Hz. Unless otherwise specified, sEPSPs were detected and captured using an EPSP template with a 2 ms rise and an 18 ms decay with the threshold set to 5 times the noise standard deviation to minimize any chance of false-positive events (see **Figure 2**). sEPSP peaks were measured as the average amplitude across a 0.3 ms period centered on the peak of the sEPSP. For consistency across all neurons and experiments, comparisons of sEPSP attributes and holding currents were made from mean values in the final 1-minute of baseline conditions (or during the final 1-minute of exposure to cholinergic antagonists, TTX, or CNO) and during the final 1-minute of exposure to ACh (or CNO), regardless of where the peak effects of ACh or CNO were observed.

**Figure 2.**
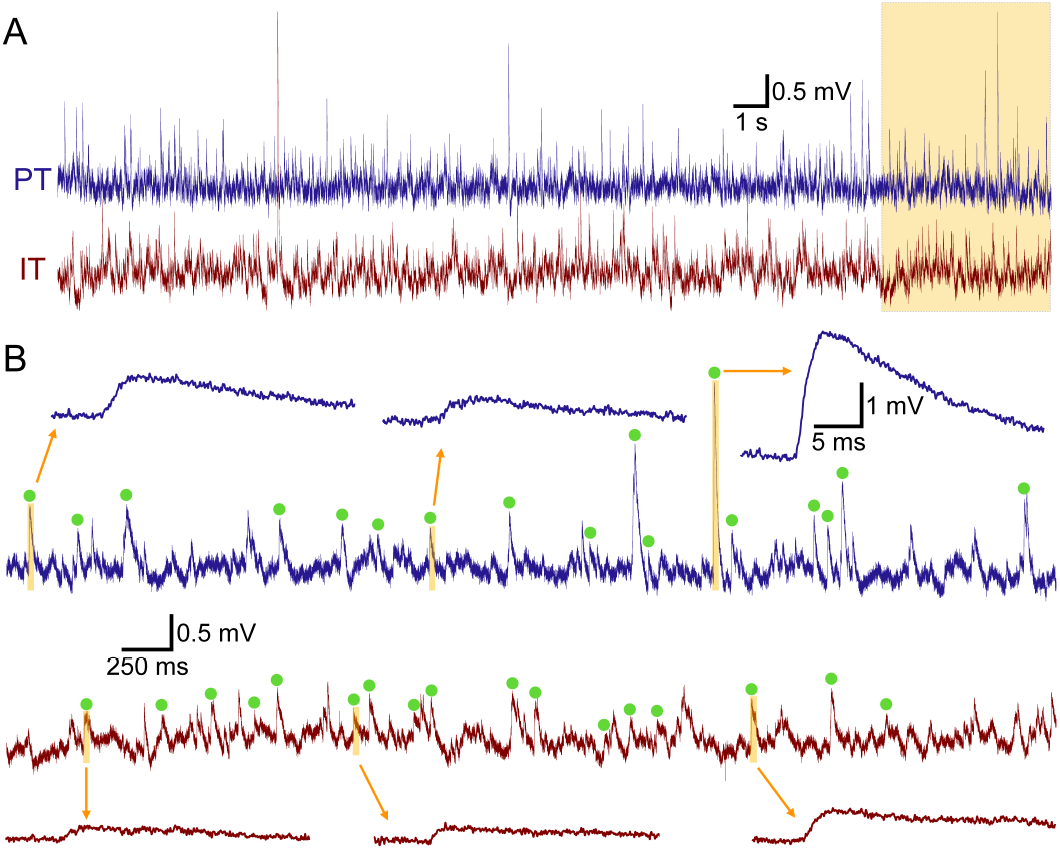
Detection of spontaneous excitatory postsynaptic potentials (sEPSPs) in IT and PT neurons. ***A***) Recordings of the membrane potentials of a pair of IT and PT neurons. The final five seconds (orange shading) are expanded in ***B. B***) Green dots show sEPSPs detected by the sEPSP template (see Methods). Several captured sEPSPs are shown at an expanded timescale.

sEPSPs in paired recordings were considered “synchronous” if their onsets occurred within an 0.5 ms window. Synchronous sEPSPs were quantified for each experimental condition (i.e., in baseline, acetylcholine, and wash conditions) as a proportion (%) of all sEPSPs occurring within the final four minutes of that condition. Proportions were averaged for each experimental pair (i.e., pair = “n”), and results compared with control *post hoc* “shuffled” pairings across all possible combinations within an experimental group. For instance, for each of the 22 PT-PT paired recordings (44 PT neurons) described below, synchronous sEPSPs were determined for all 84 shuffled pairings (42 non-paired partners for each of the two PT neurons in the experimental pair) and results averaged together to determine the “chance” level of synchronous events for each experimental pair (see **Figure 8C**). This allowed pair-wise comparison within each data set (IT-IT, IT-PT, or PT-PT) to determine whether the occurrence of synchronous sEPSPs rose above chance levels within and across experimental conditions.

**Figure 8.**
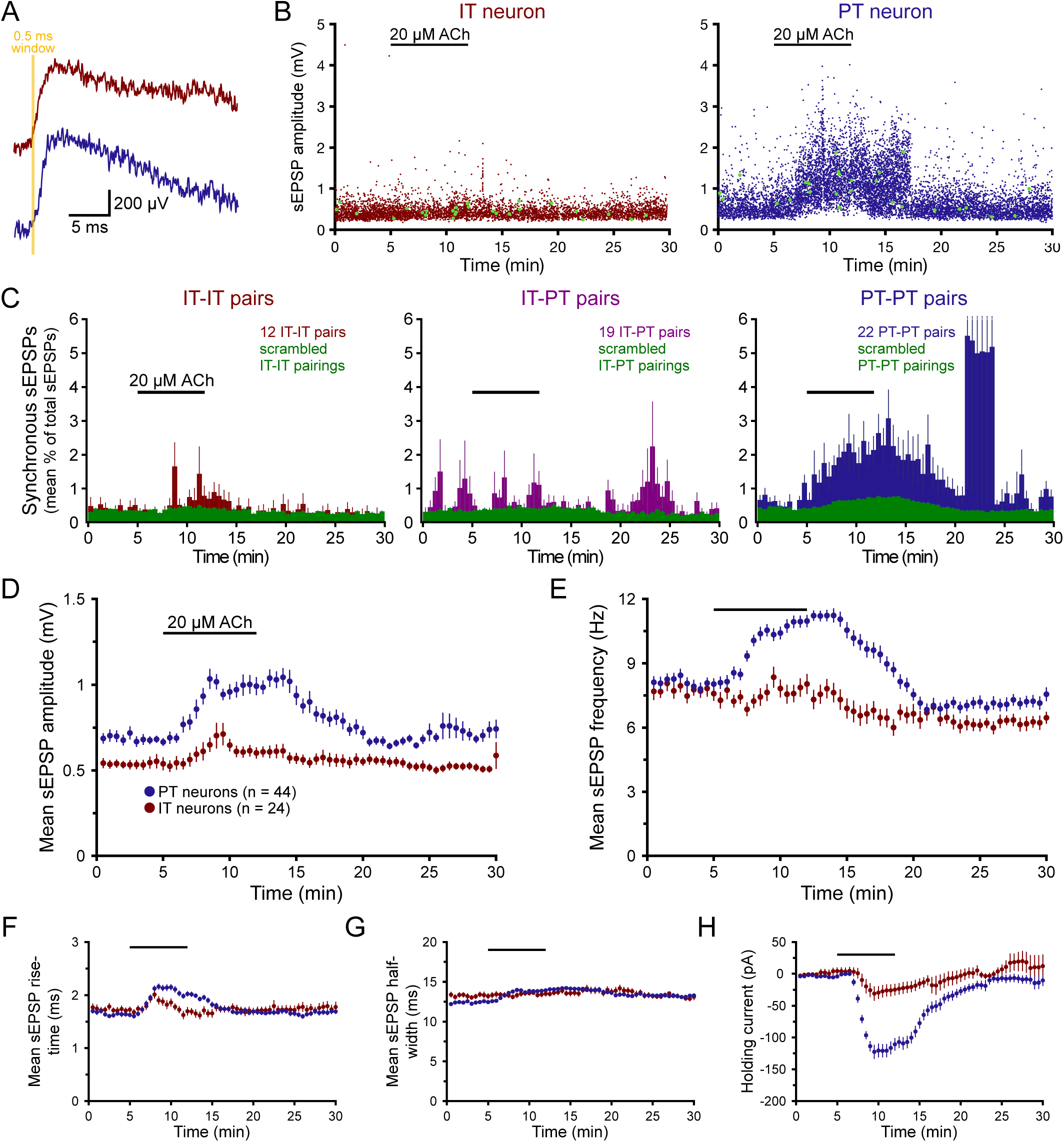
Acetylcholine (ACh) promotes synchronous excitatory input in PT-PT neuron pairs. ***A***) A pair of “synchronous” sEPSPs initiating within an 0.5 ms window (yellow) detected in a pair of IT (red trace) and PT (blue trace) neurons. ***B***) Plots of sEPSP amplitudes over 30 minutes in a paired recording of an IT (left) and PT (right) neuron. Green dots indicate synchronous EPSPs detected in both neurons. ***C***) Histograms showing the proportion (mean percentage ± SEM in 30-second bins) of all sEPSPs that occurred synchronously (onsets within an 0.5 ms window) over time for pairs of IT neurons (left; red; n = 12 pairs), IT-PT pairs (middle; purple; the same 19 pairs as shown in Figure 3), and PT-PT pairs (right; blue; n = 22 pairs). Green data indicate the proportions of sEPSPs (mean percentage ± SEM) detected as synchronous for all possible shuffled pairs to provide a measure of “chance” likelihood of synchronous events (see **Methods**). Error bars truncated during the burst of activity in a pair of PT-PT neurons (∼ minute 23). Acetylcholine increased the likelihood of synchronous events in PT-PT pairs relative to IT-IT and IT-PT pairs. ***D***) Plots of mean sEPSP amplitudes (± SEM; 30-second means) over time for the 24 IT (red) and 44 PT (blue) neurons recorded in the homotypic pairs that contributed to panel ***C***. ***E***) Mean sEPSP frequencies over time for IT and PT neurons. ***F*** - ***H***) Plots of mean sEPSP rise-times (***F***), widths at half-amplitude (***G***), and the bias currents necessary to keep neurons at their resting membrane potentials (***H***).

### Drugs

Most drugs were dissolved in water to make stock solutions (typically 10,000x concentrations) that were frozen as aliquots and used as needed during experiments. Acetylcholine (Fisher Scientific) was bath-applied with physostigmine hemisulfate (eserine; Tocris Bioscience), a blocker of acetylcholinesterase. Atropine and pirenzepine dihydrochloride were purchased from Sigma-Aldrich. Tetrodotoxin (TTX) citrate and CGP 52432 were purchased from HelloBio, Inc. Clozapine *N*-oxide (CNO) dihydrochloride was purchased from Tocris Biosciences. SR-95531 (gabazine) was purchased from MedChemExpress and dissolved in DMSO for stock solutions.

### Experimental Design and Statistical Analysis

When data are descriptive and convey variability within populations (e.g., physiological properties of IT and PT neurons), data are presented as means ± standard deviations (SD). When data are used to compare mean values (e.g., in plots of sEPSP attributes) data are presented as means ± standard error of the mean (SEM). Within group mean effects of drug treatments (e.g., ACh) and comparisons of effects in pairs of neurons (e.g., in IT-PT paired recordings) used two-tailed Student’s t-tests for paired samples. Comparisons of effects across unpaired groups utilized two-tailed Student’s t-tests for samples with unequal variance. Because of the limited usefulness of *p*-values from statistical tests (e.g., Nieuwenhuis et al., 2011; e.g., Wasserstein et al., 2019), no assumptions are made of *p*-value “significance” (i.e., ⍺). Instead, to better quantify the meaningfulness of physiological differences observed in IT and PT neurons, and the impact of ACh on sEPSP attributes, effect sizes are reported as Cohen’s *d*, a measure of the standardized mean difference between groups or conditions, according to the following formula:

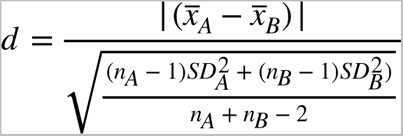

where 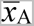 and 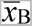 are the mean values of a parameter (e.g., sEPSP amplitude), *SD*_A_ and *SD*_B_ are the standard deviations for those values measured across groups (e.g., IT neurons vs PT neurons) or conditions (e.g., baseline conditions and in the presence of ACh), and *n*_A_ and *n*_B_ are the number of neurons in each group. Comparisons of sEPSP properties were made during the final one-minute periods of baseline conditions and ACh exposure. By convention, effect sizes ∼0.8 or larger are considered as having “large” impact (i.e., with an overlap of standard deviations being only ∼50% or less), and values of ∼0.5 and ∼0.2 considered as having “medium” or “small” impacts, respectively, according to the degree of overlap in their distributions (Cohen, 1988).

## Results

### ACh enhances spontaneous EPSPs preferentially in PT neurons

As previously described in this laboratory and others (Dembrow et al., 2010; Joshi et al., 2016; Baker et al., 2018b), ACh preferentially enhances the intrinsic excitability of neocortical PT neurons relative to neighboring IT neurons to promote corticofugal output to the thalamus and brainstem. What is less established is how ACh regulates the afferent excitatory drive of these two projection neuron subtypes. To quantify the net impact of ACh on overall excitatory drive, wholecell recordings were made from pairs of layer 5 neurons in the mouse prelimbic cortex. IT and PT neurons were classified based on their physiological responses to depolarizing and hyperpolarizing current steps (see **Methods**, **Figure 1A-D**, and **Table 1**). Identified IT and PT neurons also differed in additional physiological measures, including their resting membrane potentials, the amplitudes of afterhyperpolarizations, and action potential (AP) attributes, including spike thresholds, peaks, amplitudes, and half-widths (**Figure 1E** and **Table 2**). For both IT and PT populations, physiological attributes were similar in neurons from female (n = 78 neurons) and male (n = 128 neurons) mice.

Based on previous work showing that ACh suppresses glutamate release at some corticocortical synapses in the neocortex (Vidal and Changeux, 1993; Gil et al., 1997; Tsodyks and Markram, 1997; Hsieh et al., 2000; Atzori et al., 2005; Levy et al., 2006), it was hypothesized that exposure to ACh would reduce the amplitude and frequency of sEPSPs in IT and PT neurons in the prelimbic cortex. To test this, sEPSPs were monitored continuously in 19 pairs of layer 5 IT (mean PI of 0.74 ± 0.64 [SD]) and PT (mean PI of 18.5 ± 11.7) neurons in the mouse prelimbic cortex over a 30-minute period during which ACh (20 µM, with 10 µM eserine, a blocker of acetylcholinesterase) was bath applied for seven minutes (**Figure 3A**). Contrary to expectations, ACh *increased* sEPSP amplitudes and frequencies preferentially in PT neurons (**Figure 3A - E**; **Table 3**). Despite their lower input resistances, mean sEPSP amplitudes tended to be larger in PT neurons relative to IT neurons regardless of experimental condition (**Table 3**), whereas other sEPSP attributes, including mean frequencies, rise-times, and half-widths, were similar among the two neuron subtypes in baseline conditions (**Table 3**).

**Figure 3.**
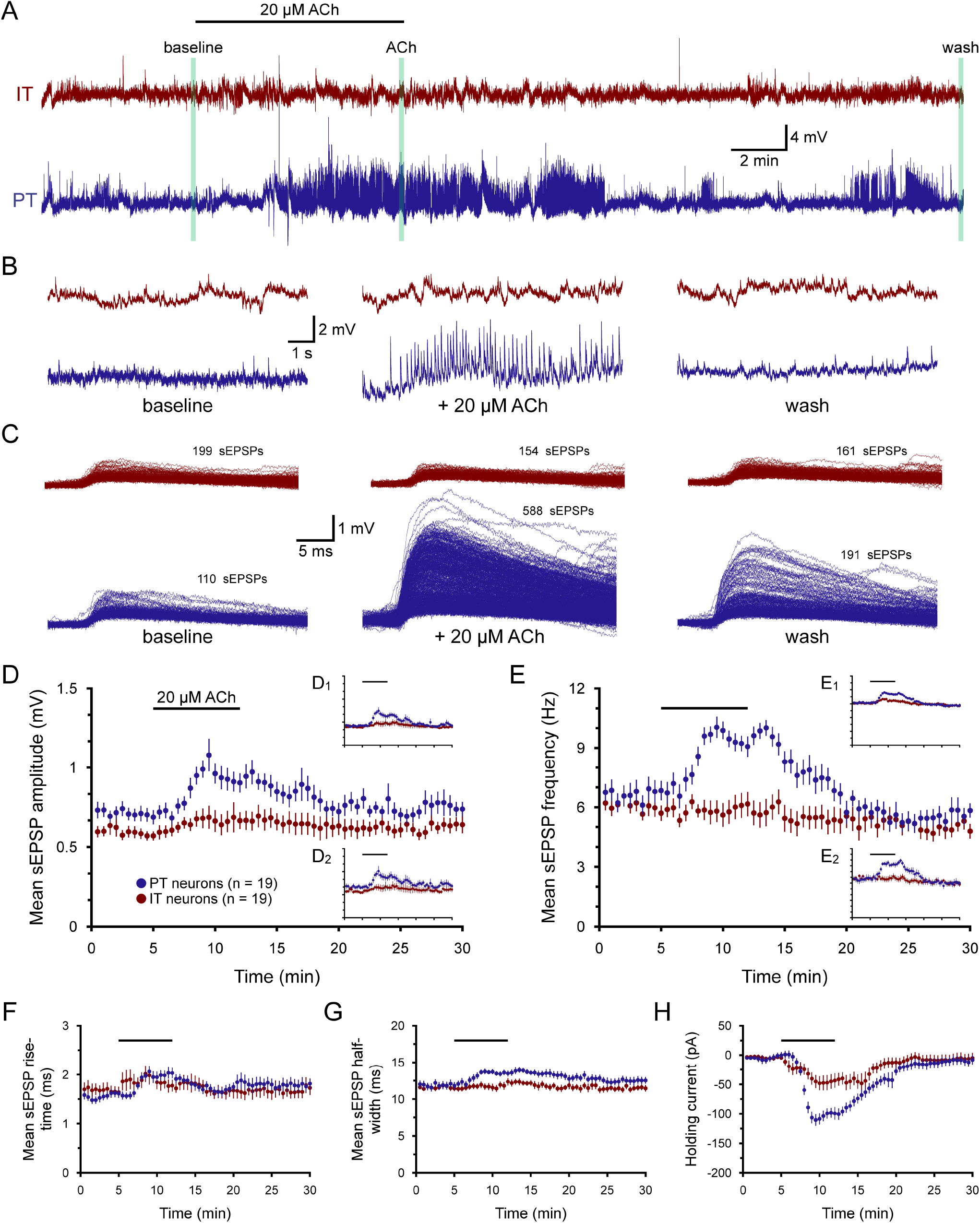
Acetylcholine (ACh) selectively enhances excitatory synaptic input to PT neurons. ***A***) In a pair of IT (red) and PT (blue) neurons, membrane potentials were recorded at the soma for thirty minutes. After a five-minute baseline period, 20 µM ACh (with 10 µM eserine) was bath applied for seven minutes. An automated slow bias (“holding”) current was used to keep neurons near their resting membrane potential throughout recordings. ***B***) Expanded 10-second traces of membrane potentials taken at the end of the baseline period (left), after seven minutes of ACh exposure (middle), and at the end of the experiment (right), as indicated in ***A***. ***C***) Superposition of all sEPSPs detected in the final 1-minute periods of baseline recording (left), ACh exposure (middle), and wash (right) in the IT (red) and PT (blue) neurons. ***D***) Plots of mean sEPSP amplitudes (± SEM) over time for 19 pairs of simultaneously recorded IT (red) and PT (blue) neurons (30-second means). Insets show the same data with EPSPs detected at 2.5x standard deviation of noise (**D_1_**) or with an EPSP template with 50% slower kinetics (**D_2_**_)_. ***E***) Similar to ***D***, including insets, but plotting mean sEPSP frequencies over time for 19 pairs of IT and PT neurons. ***F*** - ***H***) Plots of mean sEPSP rise-times (***F***), widths at half-amplitude (***G***), and the bias currents necessary to keep neurons at their resting membrane potentials (***H***).

**Table 3.** **Effects of acetylcholine on sEPSPs in IT-PT pairs (n = 19)**

| Measurement | Neuron type | Baseline | +ACh | Wash | Change in ACh | ACh vs baseline* (p-value) | Effect size (Cohen's d) |
| --- | --- | --- | --- | --- | --- | --- | --- |
| sEPSP amplitude (mV) | IT | 0.57 ± 0.03 | 0.68 ± 0.08 | 0.64 ± 0.05 | +16 ± 6% | 0.056 | 0.44 |
|  | PT | 0.69 ± 0.04 | 0.91 ± 0.06 | 0.74 ± 0.04 | +38 ± 13% | **0.007** | **0.94** |
|  | IT-PT t-test* (p) | 0.013 | 0.020 | 0.063 | 0.157 |  |  |
|  | Effect size (d) | **0.79** | **0.80** | 0.48 | 0.50 |  |  |
| sEPSP frequency (Hz) | IT | 5.8 ± 0.4 | 6.1 ± 0.6 | 5.0 ± 0.4 | +5 ± 6% | 0.435 | 0.14 |
|  | PT | 6.8 ± 0.5 | 9.2 ± 0.5 | 5.7 ± 0.4 | +50 ± 14% | <0.001 | **1.13** |
|  | IT-PT t-test* (p) | 0.079 | <0.001 | 0.223 | 0.015 |  |  |
|  | Effect size (d) | 0.49 | **1.39** | 0.40 | **0.94** |  |  |
| sEPSP rise-time (ms) | IT | 1.65 ± 0.11 | 1.77 ± 0.13 | 1.75 ± 0.15 | +10 ± 6% | 0.249 | 0.24 |
|  | PT | 1.62 ± 0.08 | 2.03 ± 0.11 | 1.79 ± 0.11 | +29 ± 7% | <0.001 | **1.02** |
|  | IT-PT t-test* (p) | 0.820 | 0.109 | 0.864 | 0.039 |  |  |
|  | Effect size (d) | 0.07 | 0.52 | 0.05 | 0.68 |  |  |
| sEPSP half-width (ms) | IT | 11.5 ± 0.4 | 12.1 ± 0.5 | 11.5 ± 0.4 | +6 ± 3% | 0.051 | 0.33 |
|  | PT | 11.9 ± 0.5 | 13.5 ± 0.5 | 12.5 ± 0.5 | +16 ± 5% | 0.003 | **0.80** |
|  | IT-PT t-test* (p) | 0.394 | 0.027 | 0.140 | 0.056 |  |  |
|  | Effect size (d) | 0.24 | **0.79** | 0.54 | 0.62 |  |  |
| Holding current (pA) | IT | -4 ± 3 | -44 ± 12 | -6 ± 8 | -40 ± 12 pA | 0.004 | **1.02** |
|  | PT | -1 ± 5 | -99 ± 9 | -10 ± 7 | -98 ± 8 pA | <0.001 | **3.03** |
|  | IT-PT t-test* (p) | 0.605 | <0.001 | 0.355 | <0.001 |  |  |
|  | Effect size (d) | 0.18 | **1.15** | 0.13 | **1.28** |  |  |

Data are shown as means ± SEM. \*Student's t-test for paired samples. Large effect sizes ( $\geq \sim 0.80$ ) are shown in bold.

Population time-courses of mean sEPSP attributes and holding currents are shown in **Figure 3D - H** for 19 IT and PT neuron pairs. The effects of ACh peaked about 5 minutes into the 7 minute exposure and typically plateaued or slightly decreased in magnitude during the final two minutes in ACh before returning to near-baseline values by the end of the 30-minute recording. For example, peak increases in sEPSP amplitudes and frequencies occurred, on average, after 4.9 ± 0.6 minutes and 5.1 ± 0.5 minutes, respectively, of ACh exposure in IT neurons, and at 4.3 ± 0.9 minutes and 5.4 ± 0.5 minutes, respectively, in PT neurons.

For the 19 pairs of IT and PT neurons tested, ACh reversibly increased sEPSP amplitudes by 16 ± 6% (mean + SEM) and 38 ± 13% in IT and PT neurons, respectively, and reversibly increased sEPSP frequencies selectively in PT neurons by 50 ± 14% (**Table 3**). The preferential impact of ACh on sEPSPs in PT neurons was not dependent on sEPSP detection threshold (**Figure 3D_1_**) or the kinetics of the EPSP template used to detect sEPSPs (**Figure 3D_2_**). ACh also preferentially increased sEPSP rise-times and half-widths in PT neurons, by 29 ± 7% and 16 ± 5%, respectively, compared to much smaller changes in IT neurons (**Figure 3F**,**G** and **Table 3**). Finally, ACh generated greater depolarizing currents in PT neurons, as reflected in the larger holding currents (-98 ± 8 pA vs -40 ± 12 pA for IT neurons) necessary to keep PT neurons at their initial resting membrane potentials during ACh exposure (**Figure 3H** and **Table 3**). This preferential postsynaptic action of ACh on PT neuron excitability is consistent with previous reports (Dembrow et al., 2010; Joshi et al., 2016; Baker et al., 2018b) showing that ACh preferentially excites PT neurons. The data presented here go further to demonstrate that ACh also preferentially increases the excitatory synaptic drive of PT neurons, suggesting that ACh will promote corticofugal output from the cortex via parallel enhancement of synaptic excitation and increased postsynaptic gain.

### Cholinergic enhancement of sEPSPs requires M1-type muscarinic receptors

In the mPFC, most postsynaptic effects of ACh in layer 5 pyramidal neurons are mediated by muscarinic ACh receptors (mAChRs; Gulledge et al., 2009; Hedrick and Waters, 2015). The test whether mAChRs are involved in regulating the excitatory drive of PT neurons, atropine (1 µM), a potent mAChR antagonist, was bath applied for five minutes prior to additional application of ACh (20 µM, with eserine; **Figure 4**). Atropine on its own had little impact on sEPSP attributes or holding currents (**Table 4**), but its presence blocked cholinergic modulation of sEPSPs and limited the amount of postsynaptic holding current necessary to keep neurons at their initial resting membrane potentials (n = 10 PT neurons; **Figure 4**, **Table 4**). These data demonstrate that cholinergic enhancement of sEPSPs in PT neurons is mediated by mAChRs.

**Figure 4.**
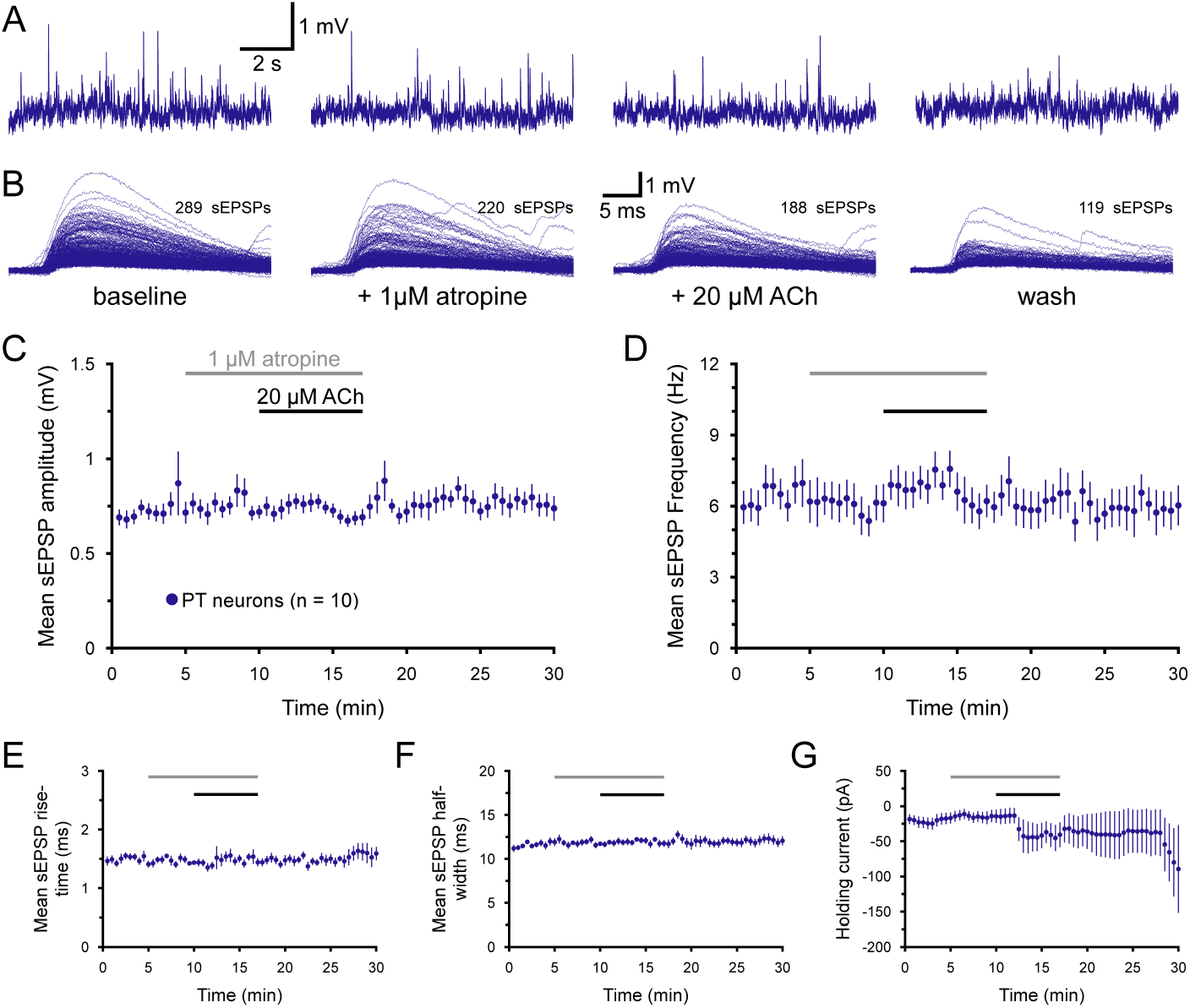
Cholinergic enhancement of sEPSPs in PT neurons is mediated by muscarinic acetylcholine (ACh) receptors. ***A***) Ten-second-long samples of membrane potential from a PT neuron taken at the end of the baseline recording period (far left), after five minutes of exposure to 1 µM atropine (a muscarinic receptor antagonist; middle left), after an additional seven minutes of exposure to atropine and 20 µM ACh (with 10 µM eserine; middle right), and after thirteen minutes of wash (far right), as indicated in ***B***. ***B***) Superimposed sEPSPs detected in the final one-minute periods of baseline recording, atropine-only exposure, atropine+ACh exposure, and wash, as indicated, for the neuron shown in ***A***. ***C*** - ***D***) Plots of mean sEPSP amplitude (***C***) and frequency (***D***) over time for 10 PT neurons exposed to atropine and ACh (30-second means). ***E*** - ***G***) Plots of mean sEPSP rise-time (***E***), width at half-amplitude (***F***), and the bias current necessary to keep neurons at their resting membrane potentials (***G***).

**Table 4.** Effects of acetylcholine on sEPSPs in the presence of atropine (n = 10 PT neurons) Measurement baseline +atropine +ACh Wash Change in ACh ACh vs.

| Measurement | baseline | +atropine | +ACh | Wash | Change in ACh | ACh vs atropine* (p-value) | Effect size (Cohen's d) |
| --- | --- | --- | --- | --- | --- | --- | --- |
| sEPSP amplitude (mV) | $0.79 \pm 0.12$ | $0.72 \pm 0.04$ | $0.69 \pm 0.04$ | $0.75 \pm 0.06$ | $-3 \pm 4\%$ | 0.257 | 0.24 |
| sEPSP frequency (Hz) | $6.6 \pm 0.9$ | $6.1 \pm 0.8$ | $6.0 \pm 0.9$ | $5.9 \pm 0.8$ | $+2 \pm 10\%$ | 0.378 | 0.12 |
| sEPSP rise-time (ms) | $1.48 \pm 0.08$ | $1.43 \pm 0.03$ | $1.50 \pm 0.07$ | $1.56 \pm 0.14$ | $+5 \pm 6\%$ | 0.424 | 0.39 |
| sEPSP half-width (ms) | $11.9 \pm 0.4$ | $11.6 \pm 0.2$ | $11.8 \pm 0.3$ | $12.0 \pm 0.5$ | $+2 \pm 3\%$ | 0.602 | 0.19 |
| Holding current (pA) | $-17 \pm 8$ | $-14 \pm 9$ | $-43 \pm 18$ | $-85 \pm 54$ | $-29 \pm 17$ pA | 0.119 | 0.64 |

Data are shown as means ± SEM. \*Student's t-test for paired samples.

It was previously reported that activation of M1-type mAChRs increase the frequency and amplitude of excitatory postsynaptic currents (EPSCs) in layer 5 pyramidal neurons in the rodent prefrontal cortex (Shirey et al., 2009). To determine whether the cholinergic enhancement of sEPSPs in PT neurons depends on M1 receptors, ACh was applied in the presence of the M1-selective antagonist pirenzepine (PZP; 1 µM). As was the case for atropine, PZP on its own had little impact on sEPSP attributes or holding currents (n = 13 PT neurons; **Figure 5**, **Table 5**). However, PZP blocked the ability of ACh to enhance sEPSP attributes and reduced ACh-induced holding currents relative to the 19 control PT neurons from **Figure 3** (*p* < 0.001; *d* = 2.1). Instead, ACh in the presence of PZP *reduced* the frequency of sEPSPs by 22 ± 9% (from 7.7 ± 0.5 Hz to 5.8 ± 0.5 Hz; **Table 5**). This suggests that ACh acting on non-M1 receptors can reduce sEPSP frequencies in PT neurons, an effect that is normally masked by stronger M1-mediated enhancement of sEPSPs.

**Figure 5.**
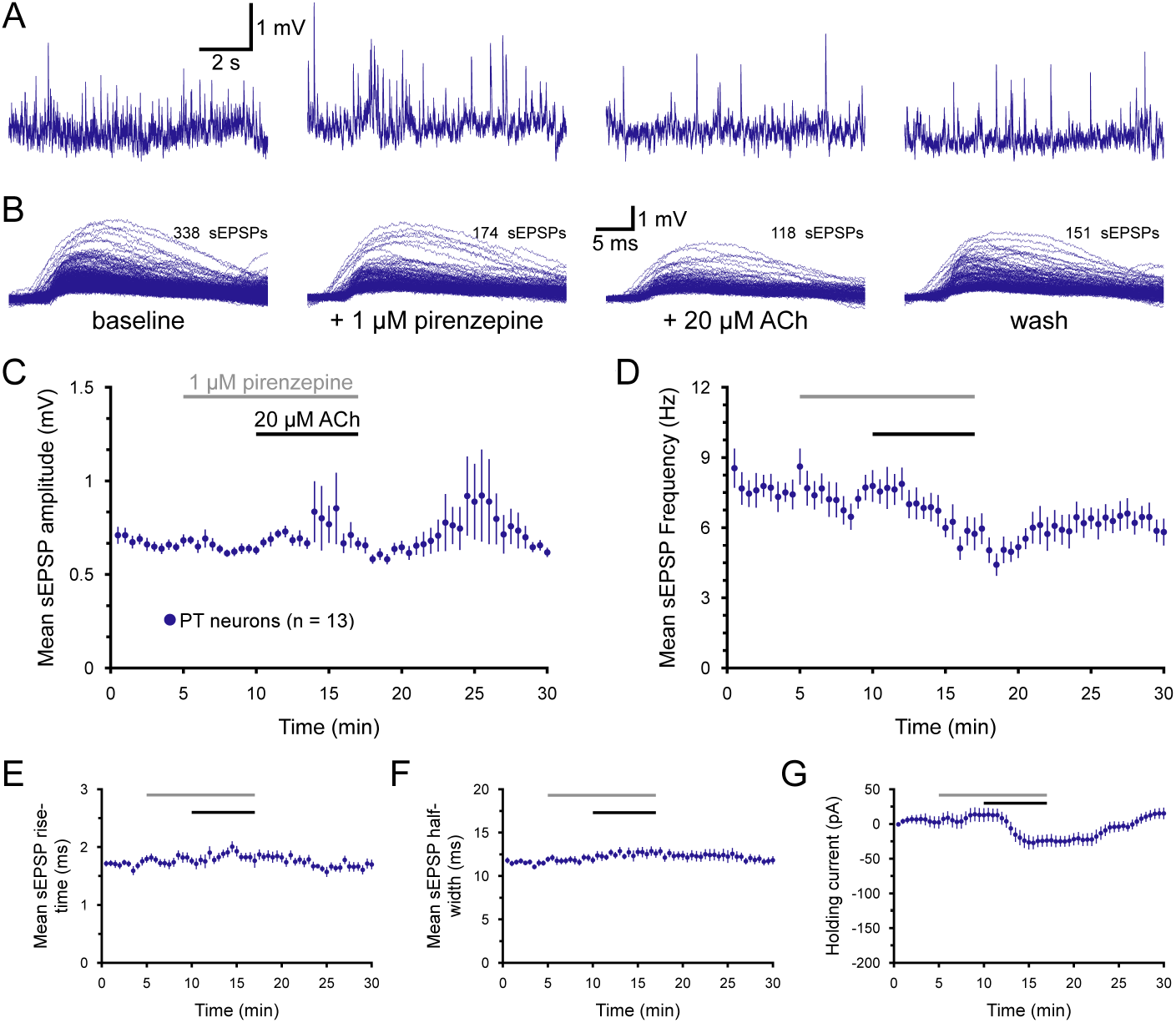
Cholinergic enhancement of sEP-SPs in PT neurons is mediated by M1-type muscarinic acetylcholine (ACh) receptors. ***A***) Ten-second-long samples of membrane po-tential from a PT neuron taken at the end of the baseline recording period (far left), after five minutes of exposure to 1 μM piren-zepine (an antagonist for M1 muscarinic receptors; middle left), after an additional seven minutes of exposure to pirenzepine and 20 μM ACh (with 10 μM eserine; middle right), and after thirteen minutes of wash (far right), as indicated in ***B***. Superimposed sEP-SPs detected in the final one-minute periods of baseline recording, pirenzepine-only ex-posure, pirenzepine+ACh exposure, and wash for the neuron shown in ***A***. ***C*** - ***D***) Plots of mean sEPSP amplitude (***C***) and frequency (***D***) over time for 13 PT neurons exposed to pirenzepine and ACh (30-second means). Note that, in the presence of pirenzepine, ACh *reduced*, rather than enhanced, sEPSP frequency in PT neurons. ***E*** - ***G***) Plots of mean sEPSP rise-time (***E***), width at half-amplitude (***F***), and the bias current necessary to keep neurons at their resting membrane potentials (***G***).

**Table 5.** Effects of acetylcholine on sEPSPs in the presence of pirenzepine (PZP; n = 13 PT neurons)

| Measurement | Baseline | +PZP | +ACh | Wash | Change in ACh | ACh vs PZP* ($p$-value) | Effect size (Cohen's $d$) |
| --- | --- | --- | --- | --- | --- | --- | --- |
| sEPSP amplitude (mV) | $0.66 \pm 0.02$ | $0.63 \pm 0.02$ | $0.69 \pm 0.05$ | $0.64 \pm 0.02$ | $+10 \pm 9\%$ | 0.328 | 0.41 |
| sEPSP frequency (Hz) | $8.0 \pm 0.6$ | $7.7 \pm 0.5$ | $5.8 \pm 0.5$ | $5.8 \pm 0.5$ | $-22 \pm 9\%$ | 0.011 | **1.05** |
| sEPSP rise-time (ms) | $1.78 \pm 0.06$ | $1.79 \pm 0.07$ | $1.80 \pm 0.10$ | $1.71 \pm 0.08$ | $0 \pm 5\%$ | 0.959 | 0.01 |
| sEPSP half-width (ms) | $11.7 \pm 0.2$ | $12.0 \pm 0.3$ | $12.7 \pm 0.4$ | $11.8 \pm 0.4$ | $+7 \pm 3\%$ | 0.074 | 0.57 |
| Holding current (pA) | $+2 \pm 9$ | $-13 \pm 10$ | $-24 \pm 9$ | $+15 \pm 8$ | $-37 \pm 7$ pA | $<0.001$ | **1.11** |

Data are shown as means ± SEM. \*Student's t-test for paired samples. Large effect sizes ( $\geq 0.80$ ) are shown in bold.

### Enhancement of sEPSPs by ACh does not require changes in GABAergic circuits

In the recording conditions employed, GABA-mediated synaptic responses are expected to reverse very close to the resting membrane potential (Gulledge and Stuart, 2003), and therefore likely do not contribute to measured sEPSPs. However, ACh might regulate GABAergic circuits in the cortex that influence the net excitatory drive of PT neurons (e.g., cholinergic disinhibition of excitatory circuits; Kruglikov and Rudy, 2008). To test whether ACh affects sEPSPs in PT neurons via modulation of cortical inhibition, blockers of both GABA_A_ (gabazine, 10 µM) and GABA_B_ (CGP52432 [CGP], 2.5 µM) receptors were bath applied seven minutes prior to additional application of ACh (n = 13; **Figure 6**). Gabazine and CGP had little if any effect on sEPSPs or holding currents on their own, and their presence did not block cholinergic enhancement of sEPSP attributes or postsynaptic holding currents in PT neurons (**Table 6**). In the presence of the GABA blockers, ACh enhanced sEPSP amplitudes and frequencies (by 28 ± 10% and 34 ± 10%, respectively), amounts similar to those observed in PT neurons in the absence of GABAergic antagonists in the IT-PT pairs described above (compare with **Table 3)**. Thus, cholinergic enhancement of excitatory drive to PT neurons does not require modulation of GABAergic synaptic transmission.

**Figure 6.**
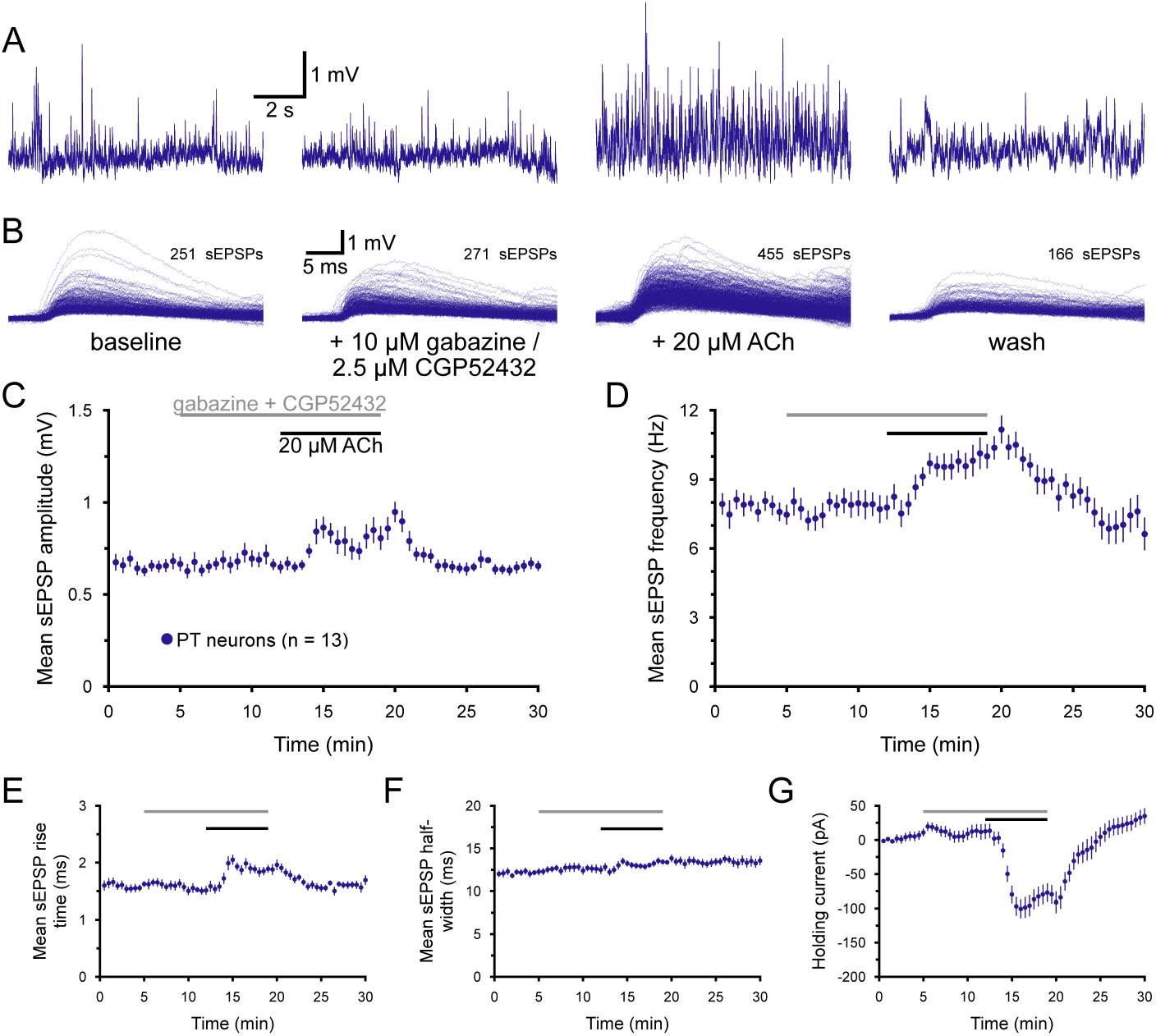
Acetylcholine (ACh) enhancement of excitatory drive does not require modula-tion of GABAergic circuits. ***A***) Ten-second-long traces of membrane potential from a PT neuron taken at the end of the baseline recording period (far left), after seven min-utes of exposure to 10 μM gabazine and 2.5 μM CGP52432 (antagonists of GABAA and GABAB receptors, respectively; middle left), after an additional seven minutes of exposure to GABA receptor blockers and 20 μM ACh (with 10 μM eserine; middle right), and after eleven minutes of wash (far right), as indicated in ***B***. Superimposed sEPSPs detected in the final one-minute periods of baseline recording, after GABA blockade, the addi-tion of ACh, and after ten minutes of wash for the neuron shown in ***A***. ***C*** - ***D***) Plots of mean sEPSP amplitude (***C***) and frequency (***D***) over time for 13 PT neurons exposed to GABA blockers and ACh (30-second means). ***E*** - ***G***) Plots of mean sEPSP rise-time (***E***), width at half-amplitude (***F***), and the bias current necessary to keep neurons at their resting membrane potentials (***G***).

**Table 6.**
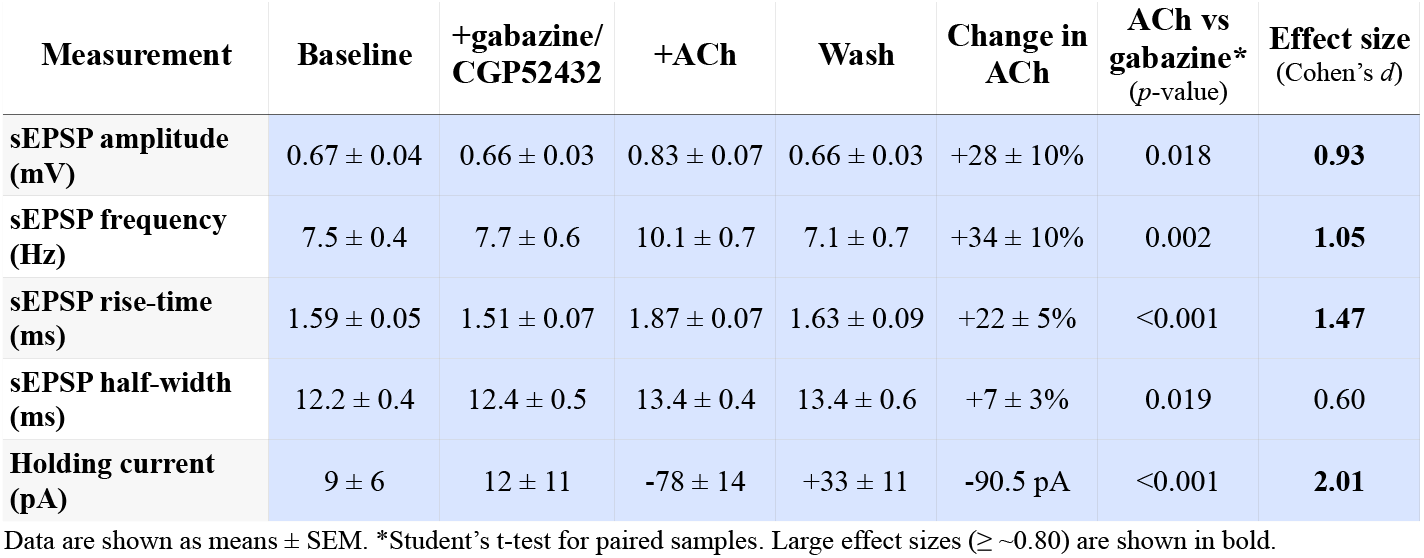
Effects of acetylcholine on sEPSPs in the absence of GABAergic transmission (n = 13 PT neurons)

| Measurement | Baseline | +gabazine/<br>CGP52432 | +ACh | Wash | Change in<br>ACh | ACh vs<br>gabazine*<br>($p$-value) | Effect size<br>(Cohen's $d$) |
| --- | --- | --- | --- | --- | --- | --- | --- |
| sEPSP amplitude<br>(mV) | $0.67 \pm 0.04$ | $0.66 \pm 0.03$ | $0.83 \pm 0.07$ | $0.66 \pm 0.03$ | $+28 \pm 10\%$ | 0.018 | **0.93** |
| sEPSP frequency<br>(Hz) | $7.5 \pm 0.4$ | $7.7 \pm 0.6$ | $10.1 \pm 0.7$ | $7.1 \pm 0.7$ | $+34 \pm 10\%$ | 0.002 | **1.05** |
| sEPSP rise-time<br>(ms) | $1.59 \pm 0.05$ | $1.51 \pm 0.07$ | $1.87 \pm 0.07$ | $1.63 \pm 0.09$ | $+22 \pm 5\%$ | $<0.001$ | **1.47** |
| sEPSP half-width<br>(ms) | $12.2 \pm 0.4$ | $12.4 \pm 0.5$ | $13.4 \pm 0.4$ | $13.4 \pm 0.6$ | $+7 \pm 3\%$ | 0.019 | 0.60 |
| Holding current<br>(pA) | $9 \pm 6$ | $12 \pm 11$ | $-78 \pm 14$ | $+33 \pm 11$ | $-90.5$ pA | $<0.001$ | **2.01** |

Data are shown as means ± SEM. \*Student's $t$ -test for paired samples. Large effect sizes ( $\geq 0.80$ ) are shown in bold.

### ACh promotes action-potential-dependent excitatory transmission in PT neurons

The “spontaneous” EPSPs described above comprise two distinct types of synaptic release. Some sEP-SPs result from synaptic release following action-potentials occurring spontaneously (i.e., not experimentally evoked) in the local network, while other sEPSPs reflect action-potential-independent “miniature” events occurring stochastically at individual synaptic boutons. To test whether ACh regulates one type of synaptic release or the other, tetrodotoxin (TTX; 1 µM), a blocker of voltage-gated sodium channels, was bath applied for seven minutes prior to the additional application of ACh (n = 13; **Figure 7**). On it’s own, TTX reduced sEPSP amplitudes (by an average of 14.9 ± 4.2%; n = 16 PT neurons; *p* = 0.028, *d* = 0.76), sEPSP frequencies (by 26 ± 3%; *p* < 0.001, *d* = 1.10), and half-widths (by 6.3 ± 1.6%; *p* = 0.002, *d* = 0.96), but had little, if any, effect on sEPSP rise times (*d* = 0.01) or holding currents (*d* = 0.08). This suggests that roughly 25% of sEPSPs recorded in baseline conditions reflect action-potential dependent release at synapses that may be somewhat larger than the average miniature EPSPs (mEPSPs; i.e., those recorded in the presence of TTX).

**Figure 7.**
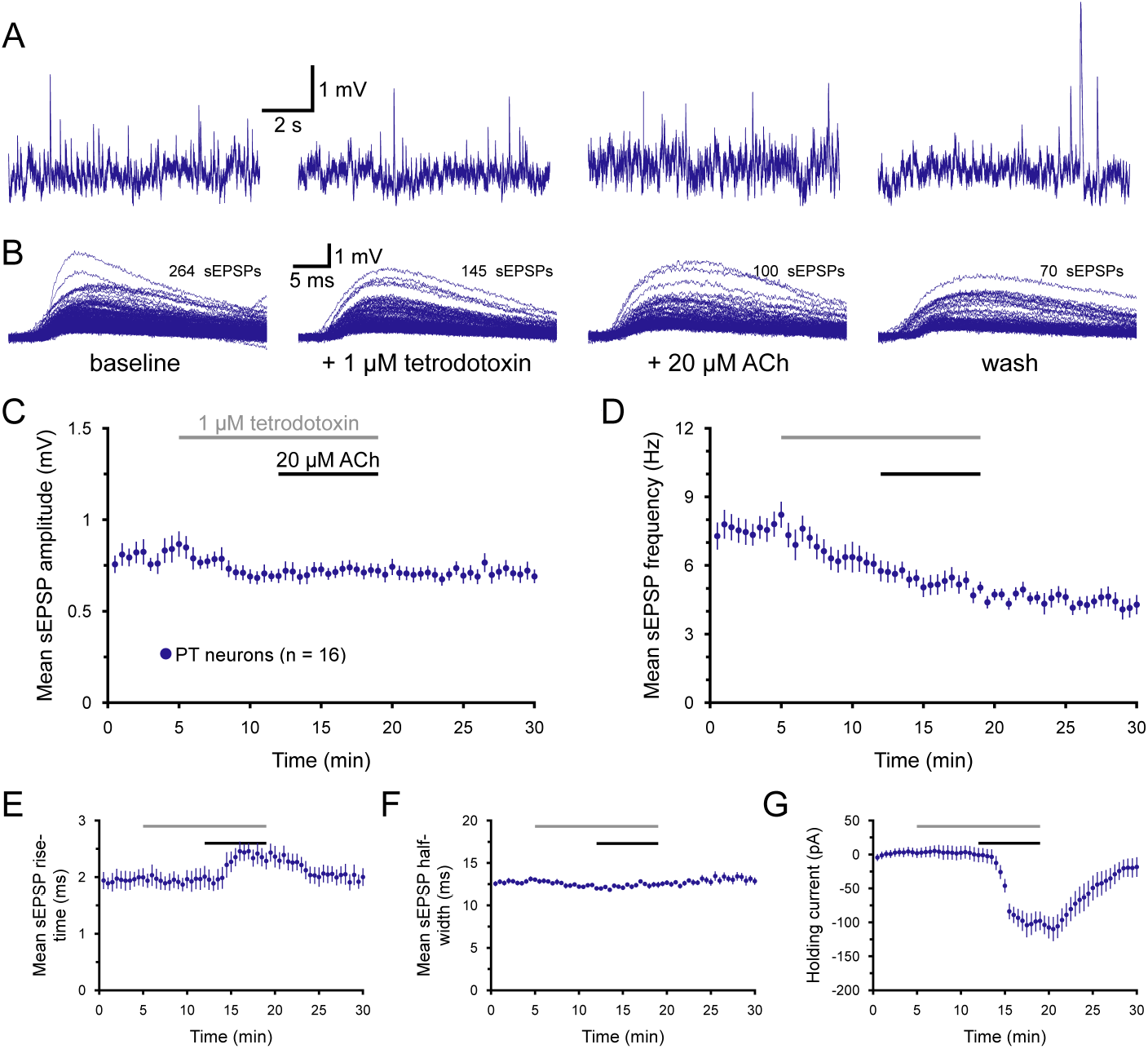
Cholinergic enhancement of excita-tory synaptic drive requires action potentials. ***A***) Ten-second-long samples of membrane potential from a PT neuron taken at the end of the baseline recording period, after seven minutes of exposure to 1 μM tetrodotoxin (TTX; a blocker of voltage-gated sodium channels), after an additional seven minutes of exposure to TTX and 20 μM acetylcholine (ACh; with 10 μM eserine), and after ten minutes of wash, as indicated in ***B***. Superim-posed sEPSPs detected in the final one-minute periods of baseline recording, TTX application, the additional application of ACh, and after ten minutes of wash for the neuron shown in ***A***. ***C*** - ***D***) Plots of mean sEPSP am-plitude (***C***) and frequency (***D***) over time for 16 PT neurons exposed to TTX and ACh (30- second bins). ***E*** - ***G***) Plots of mean sEPSP rise-time (***E***), width at half-amplitude (***F***), and the bias current necessary to keep neurons at their resting membrane potentials (***G***). Note that, in the presence of TTX, the postsynaptic effect of ACh, as quantified by changes in holding current, remains robust. The ACh-induced increase in mEPSP rise-time in the absence of other changes suggests that this effect is also mediated by postsynaptic effects of ACh on electrotonic structure.

More strikingly, the presence of TTX blocked cholinergic enhancement of mEPSP amplitudes, frequencies, and half-widths (**Table 7**). Instead, in the presence of TTX, mEPSP frequencies were *further reduced* by ACh, by an average of 15 ± 6% (**Figure 7D**, **Table 7**). While unable to enhance excitatory drive onto PT neurons, ACh in the presence of TTX continued to engage potent postsynaptic depolarizing currents (mean of -98 ± 10 pA) and increased mEPSP risetimes by 22 ± 6% (**Figure 7E**,**G**, **Table 7**). These data demonstrate that ACh selectively enhances action-potential-dependent excitatory synaptic transmission in PT neurons, and that ACh-induced changes in sEPSP rise-times likely reflect postsynaptic changes in electrotonic structure, rather than changes in synaptic release.

**Table 7.** Effects of acetylcholine on miniature EPSPs (n = 13 PT neurons)

| Measurement | Baseline | +TTX | +ACh | Wash | Change in ACh | ACh vs TTX* ($p$-value) | Effect size (Cohen's $d$) |
| --- | --- | --- | --- | --- | --- | --- | --- |
| sEPSP amplitude (mV) | $0.85 \pm 0.07$ | $0.69 \pm 0.03$ | $0.72 \pm 0.04$ | $0.71 \pm 0.04$ | $+5 \pm 5\%$ | 0.332 | 0.22 |
| sEPSP frequency (Hz) | $8.0 \pm 0.5$ | $5.9 \pm 0.4$ | $4.9 \pm 0.4$ | $4.2 \pm 0.4$ | $-15 \pm 6\%$ | 0.005 | 0.68 |
| sEPSP rise-time (ms) | $2.00 \pm 0.15$ | $1.99 \pm 0.19$ | $2.32 \pm 0.15$ | $1.96 \pm 0.14$ | $+22 \pm 6\%$ | 0.016 | 0.49 |
| sEPSP half-width (ms) | $13.1 \pm 0.3$ | $12.2 \pm 0.2$ | $12.4 \pm 0.3$ | $13.0 \pm 0.5$ | $+2 \pm 2\%$ | 0.260 | 0.27 |
| Holding current (pA) | $+3 \pm 7$ | $0 \pm 10$ | $-98 \pm 14$ | $-19 \pm 14$ | $-98 \pm 10$ pA | $<0.001$ | **1.96** |

Data are shown as means ± SEM. \*Student's $t$ -test for paired samples. Large effect sizes ( $\geq \sim 0.80$ ) are shown in bold.

### ACh promotes synchronous input to PT neurons

Unlike “miniature” synaptic transmission, which occurs stochastically at individual boutons, action-potential-dependent transmission is expected to promote transmitter release (with some probability) from all boutons on a given axon. Given that ACh promotes action-potential-dependent glutamate release, it is plausible that ACh may promote synchronous input to populations of neurons innervated by a common set of axons. To test whether this was the case in the 19 IT-PT paired recording experiments described above, a 0.5 ms window was used to find “synchronous” events occurring in both neurons (**Figure 8A, B**). The probability of synchronous events in baseline conditions was low (0.84 ± 1.45 [SD] of sEPSPs; n = 19 IT-PT pairs) but was greater than double the “chance” level of synchronous events occurring within a 0.5 ms window (0.35 ± 0.07%; n = 36 shuffled IT-PT pairings for each of 19 experimental pairs; **Figure 8C**, **Table 8**). Application of ACh failed to enhance synchronous input probabilities in IT-PT pairs (0.80 ± 1.12% of sEPSPs) and had only a small effect in shuffled IT-PT pairs (0.43 ± 0.09% in the presence of ACh; **Figure 8C**), that reflected the increase in total sEPSP occurring in ACh conditions (particularly in the PT neurons). These data demonstrate that ACh does not enhance the likelihood of synchronous excitatory input in IT and PT neurons.

**Table 8.** Effects of acetylcholine on the probability of synchronous input recorded in heterotypic and homotypic neuron pairs.

| Group | n | Mean % coincident sEPSPs |  |  | Raw change in ACh | ACh vs baseline* (p-value) | Effect size (Cohen's d) |
| --- | --- | --- | --- | --- | --- | --- | --- |
|  |  | Baseline | +ACh | Wash |  |  |  |
| IT-IT pairs | 12† | 0.37 ± 0.25 | 0.83 ± 0.85 | 0.24 ± 0.22 | +0.5 ± 1.0% | 0.125 | 0.73 |
| IT-IT shuffled | 12 (x20)# | 0.36 ± 0.04 | 0.44 ± 0.09 | 0.27 ± 0.04 | +0.1 ± 0.1% | 0.002 | **1.26** |
| Pairs vs shuffled | t-test* (p) | 0.859 | 0.141 | 0.633 | 0.205 |  |  |
|  | Effect size (d) | 0.07 | 0.64 | 0.18 | 0.55 |  |  |
| IT-PT pairs | 19† | 0.84 ± 1.45 | 0.80 ± 1.12 | 0.31 ± 0.34 | 0 ± 0.9% | 0.840 | 0.03 |
| IT-PT shuffled | 19 (x36)# | 0.35 ± 0.07 | 0.43 ± 0.09 | 0.24 ± 0.05 | +0.8 ± 0.1% | 0.002 | **0.99** |
| Pairs vs shuffled | t-test* (p) | 0.149 | 0.175 | 0.341 | 0.534 |  |  |
|  | Effect size (d) | 0.48 | 0.46 | 0.30 | 0.20 |  |  |
| PT-PT pairs | 22† | 0.60 ± 0.36 | 2.09 ± 2.11 | 0.76 ± 1.02 | +1.5 ± 2.1% | 0.003 | **0.98** |
| PT-PT shuffled | 22 (x84)# | 0.42 ± 0.05 | 0.67 ± 0.05 | 0.35 ± 0.06 | +0.3 ± 0.05% | <0.001 | **4.92** |
| Pairs vs shuffled | t-test* (p) | 0.023 | 0.005 | 0.075 | 0.011 |  |  |
|  | Effect size (d) | 0.70 | **0.95** | 0.57 | **0.85** |  |  |

Data are shown as means ± SD for the proportion (%) of coincident (occurring within a 0.5 ms window) sEPSPs relative to all sEPSPs in the final four minutes each of baseline recording, ACh exposure, and wash. \*Percentage of synchronous sEPSPs for each experimental condition were averaged for each recorded pair (i.e., pairs treated as "n"). #For each recorded pair, data from every possible shuffled combination for both neurons were averaged (see **Methods**). \*Student's t-test for paired samples. Large effect sizes ( $\geq \sim 0.80$ ) are shown in bold.

To test whether ACh might promote synchronous excitatory input in homotypic pairs of neurons, addi- tional paired recordings of sEPSPs were made from IT-IT (n = 12) and PT-PT (n = 22) pairs (**Figure 8C**, **Table 8**). In IT-IT pairs, ACh increased the probability of synchronous sEPSPs from 0.37 ± 0.25% [SD] of sEP-SPs in baseline conditions to 0.83 ± 0.85% in the final four minutes of ACh exposure. This compared to a smaller but more consistent increase in “synchronous” sEPSPs in shuffled pairs (n = 20 shuffled IT pairings per experimental pair; **Table 8**) that again reflects the Gulledge - ACh activates networks of PT neurons small increase in sEPSP frequencies observed in IT neurons exposed to ACh (see **Figure 3** and **Table 3**). Notably, while the proportions of synchronous sEPSPs in IT-IT paired recordings and shuffled IT-IT pairings were almost identical in baseline and wash conditions, in the presence of ACh the proportion of synchronous events in actual IT-IT pairs was almost double that observed in shuffled data (**Table 8**), demonstrating that, unlike in IT-PT pairs, ACh modestly (*p* = 0.125, *d* = 0.73) facilitates synchronous input in IT-IT pairs.

In PT-PT pairs (n = 22), ACh increased the proportion of synchronous sEPSPs to a much greater extent (**Figure 8C**, **Table 8**). Whereas the proportion of synchronous sEPSPs in PT-PT pairs in baseline conditions was about 50% greater than in shuffled pairings (0.60 ± 0.36% vs 0.42 ± 0.05%, respectively), the addition of ACh increased the fraction of synchronous sEPSPs in PT-PT pairs to 2.09 ± 2.11%, such that it was three times that observed in shuffled data (0.67 ± 0.05%; *p* = 0.005, *d* = 0.95). When comparing the raw change in the percent of synchronous input occurring in ACh (relative to baseline), PT-PT pairs showed larger mean changes (+1.49 ± 2.07) than observed in IT-IT (0.46 ± 0.96; *p* = 0.057, *d* = 0.58 relative to PT-PT pairs) or IT-PT (-0.04 ± 0.83; *p* < 0.004, *d* = 0.94 relative to PT-PT pairs) pairs. These data demonstrate that ACh preferentially promotes the occurrence of synchronous sEPSPs in PT neurons.

It is worth noting that, despite testing for unitary connectivity in all paired recordings in this study, no unitary connections were identified, regardless of which projection neuron subtypes were targeted. This likely reflects severed axons in coronal slices of prefrontal cortex, as in all experiments slices were placed such that apical dendrites were level with, or descended into, the slice to preserve dendritic architecture. A consequence of this is that axons of neurons close to the surface of the slice are frequently severed a few hundred µm from the soma.

In addition to measuring the impact of ACh on synchronous inputs in PT-PT neuron pairs (above), the effects of ACh on sEPSP attributes were compared across neuron subtypes in the 12 IT-IT (24 total IT neurons) and 22 PT-PT (44 total PT neurons) paired recordings described above. As was found in IT-PT pairs, ACh preferentially enhanced sEPSP amplitudes (by 53 ± 10%), frequencies (by 42 ± 6%), rise-times (by 25 ± 4%), and half-widths (by 12 ± 2%) in PT neurons (n = 44; *p* values at or below 0.005 and effect sizes ranging from 0.70 to 1.21 when compared to the 24 IT neurons from IT-IT pairs; **Figure 8D - H**; **Table 9**). Similarly, the holding currents necessary to maintain neurons at their resting membrane potentials were much larger in PT neurons (-108 ± 10 pA) than in IT neurons (-29 ± 5 pA). Thus, the results from paired IT-IT and PT-PT recordings (68 neurons in total) represent a replication of the ACh effects on sEPSPs observed in the original 19 IT-PT neuron pairs (compare **Figure 8D - H** with **Figure 3D - H**), providing additional confidence that cholinergic enhancement of sEPSPs occurs preferentially in PT neurons.

**Table 9.** Effects of acetylcholine on sEPSPs in IT and PT neurons recorded as homotypic pairs.

| Measurement | Neuron type (n) | Baseline | +ACh | Wash | Change in ACh | ACh vs baseline* (p-value) | Effect size (Cohen's d) |
| --- | --- | --- | --- | --- | --- | --- | --- |
| sEPSP amplitude (mV) | IT (24) | 0.55 ± 0.04 | 0.61 ± 0.03 | 0.55 ± 0.06 | +16 ± 6% | 0.026 | 0.35 |
| | PT (44) | 0.68 ± 0.02 | 1.01 ± 0.05 | 0.74 ± 0.05 | +53 ± 10% | <0.001 | **1.20** |
|  | IT-PT t-test† (p) | 0.006 | <0.001 | 0.005 | 0.001 |  |  |
|  | Effect size (d) | **0.82** | **1.32** | 0.65 | 0.70 |  |  |
| sEPSP frequency (Hz) | IT (24) | 7.7 ± 0.4 | 8.0 ± 0.5 | 6.3 ± 0.3 | +4 ± 5% | 0.397 | 0.12 |
| | PT (44) | 8.0 ± 0.3 | 10.9 ± 0.3 | 7.1 ± 0.3 | +42 ± 6% | <0.001 | **1.65** |
|  | IT-PT t-test† (p) | 0.497 | <0.001 | 0.050 | <0.001 |  |  |
|  | Effect size (d) | 0.18 | **1.55** | 0.47 | **1.12** |  |  |
| sEPSP rise-time (ms) | IT (24) | 1.72 ± 0.06 | 1.66 ± 0.08 | 1.75 ± 0.09 | -2 ± 4% | 0.272 | 0.17 |
| | PT (44) | 1.65 ± 0.05 | 2.02 ± 0.05 | 1.70 ± 0.04 | +25 ± 4% | <0.001 | **1.11** |
|  | IT-PT t-test† (p) | 0.399 | <0.001 | 0.603 | <0.001 |  |  |
|  | Effect size (d) | 0.21 | **1.05** | 0.15 | **1.21** |  |  |
| sEPSP half-width (ms) | IT (24) | 13.3 ± 0.4 | 13.5 ± 0.3 | 13.0 ± 0.4 | +2 ± 3% | 0.492 | 0.10 |
| | PT (44) | 12.5 ± 0.2 | 14.0 ± 0.2 | 13.4 ± 0.2 | +12 ± 2% | <0.001 | **1.09** |
|  | IT-PT t-test† (p) | 0.102 | 0.187 | 0.366 | 0.005 |  |  |
|  | Effect size (d) | 0.47 | 0.37 | 0.23 | **0.80** |  |  |
| Holding current (pA) | IT (24) | +4 ± 6 | -25 ± 9 | +11 ± 19 | -29 ± 5 pA | <0.001 | **0.78** |
| | PT (44) | -5 ± 3 | -111 ± 10 | -6 ± 9 | -108 ± 10 pA | <0.001 | **2.22** |
|  | IT-PT t-test† (p) | 0.159 | <0.001 | 0.413 | <0.001 |  |  |
|  | Effect size (d) | 0.42 | **1.45** | 0.24 | **1.46** |  |  |

Data are shown as means ± SEM. \*Student's t-tests for paired samples were used to compare sEPSP attributes across baseline and cholinergic conditions. †Student's t-tests assuming unequal variances were used for comparisons of sEPSP attributes in IT and PT neurons for each experimental condition. Large effect sizes ( $\geq \sim 0.80$ ) are shown in bold.

### Silencing PT neurons blocks cholinergic enhancement of excitatory drive

The results described above demonstrate that ACh enhances action-potential-dependent glutamate release preferentially onto PT neurons in the mouse mPFC. What might the mechanism be for this effect? A number of observations suggest that cholinergic excitation of PT neurons, as a population, could explain the selective impact of ACh on sEPSPs in those same neurons. First, the M1 receptors implicated in cholinergic enhancement of sEPSP amplitudes and frequencies also preferentially promote persistent action potential generation in PT neurons (Gulledge et al., 2009; Dembrow et al., 2010; Joshi et al., 2016; Baker et al., 2018b). Thus, M1-dependent activation of populations of synaptically coupled PT neurons could explain an increase in TTX-sensitive sEPSP frequency. Second, PT axons preferentially innervate other PT neurons, rather than IT neurons, in local cortical circuits (Morishima and Kawaguchi, 2006; Brown and Hestrin, 2009; Kiritani et al., 2012). This could explain why cholinergic enhancement of excitatory drive is preferential to PT neurons. Finally, unitary PT-PT synaptic connections tend to be larger than unitary connections in IT-IT or IT-PT pairs (Morishima and Kawaguchi, 2006; Morishima et al., 2011), which would explain the increase in sEPSP amplitude, and perhaps half-width, following ACh application.

To test whether recurrent network activity in populations of PT neurons contributes to the cholinergic increase in sEPSPs in PT neurons, a retrograde AAV virus encoding the G_i_-coupled inhibitory hM4Di receptor was injected into the pons of mice three weeks before experiments to allow targeted silencing of PT neurons (**Figure 9**). Bath application of CNO (5 µM; an agonist for hM4Di receptors) was effective in inhibiting PT neurons, as evident from the large positive holding currents (+99 ± 14 pA; n = 21 PT neurons) necessary to maintain neurons at their initial resting membrane potential (**Figure 9I**, **Table 10**).

**Figure 9.**
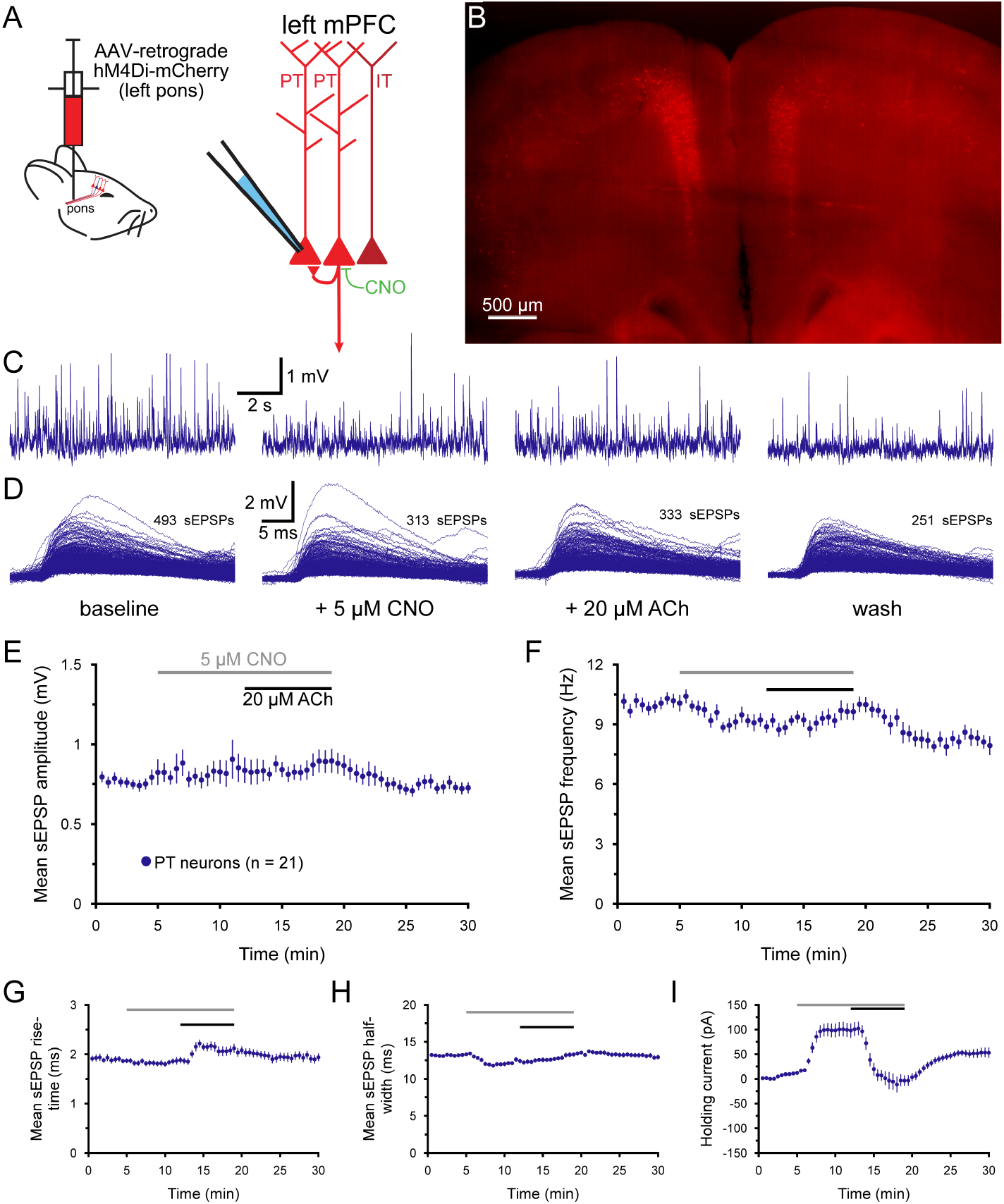
Chemogenetic inhibition of PT neurons blocks cholinergic enhancement of excitatory drive. ***A***) Diagram of injection of AAV-retrograde hM4Di-mCherry virus into the pons to express hM4Di receptors selec-tively in PT neurons in the cortex (left), and of recording setup using clozapine N-oxide (CNO; 5 μM) to silence PT neurons (right). ***B***) Image of hM4Di-m-Cherry-expressing layer 5 PT neurons in the medial prefrontal cortex (5x objective, artificially colored) three weeks after virus injection into the pons. ***C***) Ten-second-long traces of membrane poten-tial from a hM4Di-mCherry-expressing PT neuron taken at the end of the baseline record-ing period (far left), after seven minutes of exposure to 5 μM CNO (to activate hM4Di receptors; middle left), after an additional seven minutes of exposure to CNO and 20 μM acetylcholine (ACh; with 10 μM eserine; middle right), and after ten minutes of wash (far right), as indicated in ***D***. ***D***) Superim-posed sEPSPs detected in the final one-minute periods of baseline recording, after CNO application, following the additional application of ACh, and after ten minutes of wash for the neuron shown in ***C***. ***E*** - ***F***) Plots of mean sEPSP amplitude (***E***) and frequency (***F***) over time for 21 PT neurons exposed to CNO and ACh (30-second means). ***G*** - ***I***) Plots of mean sEPSP rise-time (***G***), width at half-amplitude (***H***), and the bias current nec-essary to keep neurons at their resting mem-brane potentials (***I***). Note that CNO induced a large outward current before ACh induced its own inward current, and that ACh induced an increase in sEPSP rise-times and holding currents even while having no effect on sEP-SP amplitudes, frequencies, or half-widths (compare with Figure 7C - G).

**Table 10.** Effects of acetylcholine on sEPSPs after chemogenetic silencing of PT neurons (n = 21 PT.

| Measurement | Baseline | + CNO | +ACh | Wash | Change in ACh | ACh vs CNO* (p-value) | Effect size (Cohen's d) |
| --- | --- | --- | --- | --- | --- | --- | --- |
| sEPSP amplitude (mV) | 0.81 ± 0.07 | 0.85 ± 0.08 | 0.89 ± 0.07 | 0.73 ± 0.04 | +10 ± 6% | 0.512 | 0.14 |
| sEPSP frequency (Hz) | 10.1 ± 0.4 | 9.1 ± 0.3 | 9.6 ± 0.4 | 8.0 ± 0.4 | +8 ± 5% | 0.175 | 0.37 |
| sEPSP rise-time (ms) | 1.86 ± 0.05 | 1.87 ± 0.07 | 2.10 ± 0.09 | 1.92 ± 0.08 | +13 ± 4% | 0.013 | 0.63 |
| sEPSP half-width (ms) | 13.3 ± 0.2 | 12.5 ± 0.3 | 13.3 ± 0.3 | 12.9 ± 0.3 | +7 ± 2% | 0.002 | 0.55 |
| Holding current (pA) | +11 ± 3 | +99 ± 14 | -3 ± 11 | +53 ± 10 | -102 ± 11 pA | <0.001 | **1.73** |

Data are shown as means ± SEM. \*Student's t-test for paired samples. Large effect sizes ( $\geq 0.80$ ) are shown in bold.

CNO on its own had little impact on sEPSP amplitudes or rise-times, but modestly decreased sEPSP frequencies (by 9.6 ± 3%; n = 21; *p* = 0.002, *d* = 0.72) and half-widths (by 5.8 ± 1.9%; *p* = 0.006, *d* = 0.65). In the continued presence of CNO, application of ACh activated inward currents in PT neurons (-102 ± 11 pA, returning holding currents to near zero) and increased sEPSP rise-times (by 13 ± 4%), indicating relatively normal postsynaptic actions of ACh (e.g., compare to **Figure 7E, G**). However, CNO abolished cholinergic enhancement of sEPSP amplitudes and frequencies (**Figure 9C - H**, **Table 10**), suggesting that the increases in sEPSP amplitudes and frequencies normally observed in ACh require AP generation in networks of interconnected PT neurons.

In another group of animals the retrograde AAV hM4Di virus was injected into the left mPFC to express hM4Di in IT neurons in the contralateral cortex (**Figure 10A**,**B**). Recordings of sEPSPs were then made in PT neurons in the contralateral hemisphere. Initial application of 5 µM CNO had little effect on sEPSP amplitudes or holding currents, but reduced sEPSP frequencies by 11.9 ± 4.9% (n = 12; *p* = 0.024, *d* = 0.59; **Figure 10C**-**I**; **Table 11**). Application of ACh in the continued presence of CNO enhanced sEPSP amplitudes (by 33 ± 8%), frequencies (by 46 ± 12%), rise-times, and half-widths (**Figure 10C**-**I**, **Table 11**). ACh also induced depolarizing postsynaptic currents (mean change in holding current was -114 ± 9 pA). The effects of ACh on sEPSPs in PT neurons after chemogenetic silencing of IT neurons were of similar magnitude to those observed in PT neurons in control conditions (compare results in **Table 11** with those in **Table 3**).

**Figure 10.**
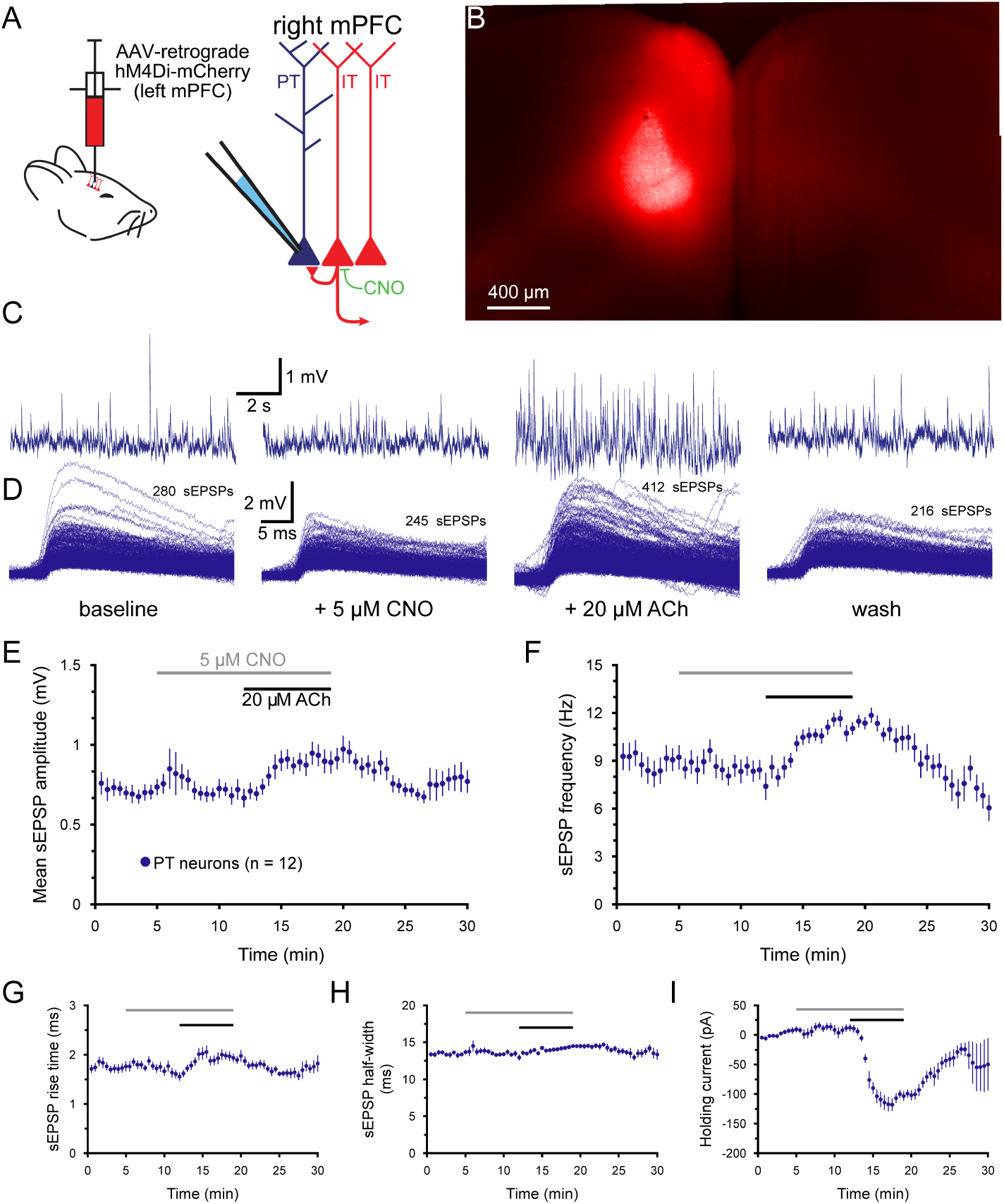
Chemogenetic inhibition of IT neurons in the medial prefrontal cortex (mPFC) does not block cholinergic enhance-ment of excitatory drive to PT neurons, ***A***) Diagram of injection of AAV-retrograde hM4Di-mCherry virus into the left mPFC (left), and of recording setup using clozapine N-oxide (CNO; 5 μM) to silence IT neurons in the contralateral hemisphere (right). ***B***) Image of hM4Di-m-Cherry-expression in the mFPC (5x objective, artificially colored) three weeks after virus injection into the left mPFC. ***C***) Ten-second-long traces of membrane potential from a hM4Di-mCherry-expressing PT neuron taken at the end of the baseline recording period (far left), after seven minutes of exposure to 5 μM CNO (to activate hM4Di receptors; middle left), after an additional seven minutes of exposure to CNO and 20 μM acetylcholine (ACh; with 10 μM eserine; middle right), and after ten minutes of wash (far right), as indicated in ***D***. ***D***) Superimposed sEPSPs detected in the final one-minute periods of baseline recording, after CNO application, following the additional application of ACh, and after ten minutes of wash for the neuron shown in ***C***. ***E*** - ***F***) Plots of mean sEPSP amplitude (***E***) and frequency (***F***) over time for 13 PT neurons exposed to CNO and ACh (30-second means). ***G*** - ***I***) Plots of mean sEPSP rise-time (***G***), width at half-amplitude (***H***), and the bias current necessary to keep neurons at their resting mem-brane potentials (***I***).

**Table 11.** Effects of acetylcholine on sEPSPs after chemogenetic silencing of IT neurons (n = 12 PT neurons)

| Measurement | Baseline | + CNO | +ACh | Wash | Change in ACh | ACh vs CNO* (p-value) | Effect size (Cohen's d) |
| --- | --- | --- | --- | --- | --- | --- | --- |
| sEPSP amplitude (mV) | 0.72 ± 0.05 | 0.69 ± 0.06 | 0.89 ± 0.07 | 0.78 ± 0.08 | +33 ± 8% | 0.057 | **0.85** |
| sEPSP frequency (Hz) | 9.1 ± 0.8 | 7.9 ± 0.6 | 10.9 ± 0.5 | 6.4 ± 0.8 | +46 ± 12% | 0.003 | **1.65** |
| sEPSP rise-time (ms) | 1.74 ± 0.10 | 1.58 ± 0.08 | 1.95 ± 0.09 | 1.79 ± 0.15 | +26 ± 7% | 0.005 | **1.23** |
| sEPSP half-width (ms) | 13.5 ± 0.3 | 13.2 ± 0.3 | 14.4 ± 0.2 | 13.4 ± 0.6 | +10 ± 2 | 0.004 | **1.44** |
| Holding current (pA) | +8 ± 5 | +12 ± 6 | -92 ± 12 | +44 ± 40 | -114 ± 9 pA | <0.001 | **4.86** |

Data are shown as means ± SEM. \*Student's t-test for paired samples. Large effect sizes ( $\geq 0.80$ ) are shown in bold.

The chemogenetic experiments described above demonstrate that network activity in PT, but not IT, populations is necessary for cholinergic enhancement of sEPSPs in PT neurons. To test whether selective excitation of PT neuron populations might be sufficient to mimic the effect of ACh on sEPSPs, a retrograde AAV encoding the G_q_-coupled DREADD hM3Dq was injected into the left pons (**Figure 11A, B**). Three weeks later, recordings of sEPSPs were made in nine hM3Dq^+^ PT neurons in the ipsilateral mPFC. Although bath application of CNO (5 µM) led to extremely large and irreversible (within 20 min) holding currents (mean of -152 ± 29 pA; **Figure 11G**, **Table 12**), CNO had little if any effect on EPSPs (**Figure 11C**-**F**, **Table 12**). The lack of effect of ACh on sEPSP rise-times, an otherwise post-synaptic effect of ACh (see **Figure 7**), suggests that CNO-induced activation of hM3Dq in PT neurons does not fully mimic M1 receptor stimulation. This may reflect differences in the subcellular localizations of DREADD and M1 receptors, or over-activation of G_q_-coupled cascades in hM3Dq-expressing neurons. For instance, broad AAV-mediated expression of hM3Dq in PT neurons, including within axons (e.g., Mitew et al., 2018; Liang et al., 2020) that are normally devoid of M1 receptors (Oda et al., 2018), could lead to depolarization block (Yi et al., 2015) and impaired synaptic transmission at PT terminals. These considerations complicate interpretation of this negative result following hM3Dq activation in PT neurons. Alternative approaches will be necessary to fully test whether selective activation of PT neurons is sufficient to mimic the effect of ACh on sEPSPs.

**Figure 11.**
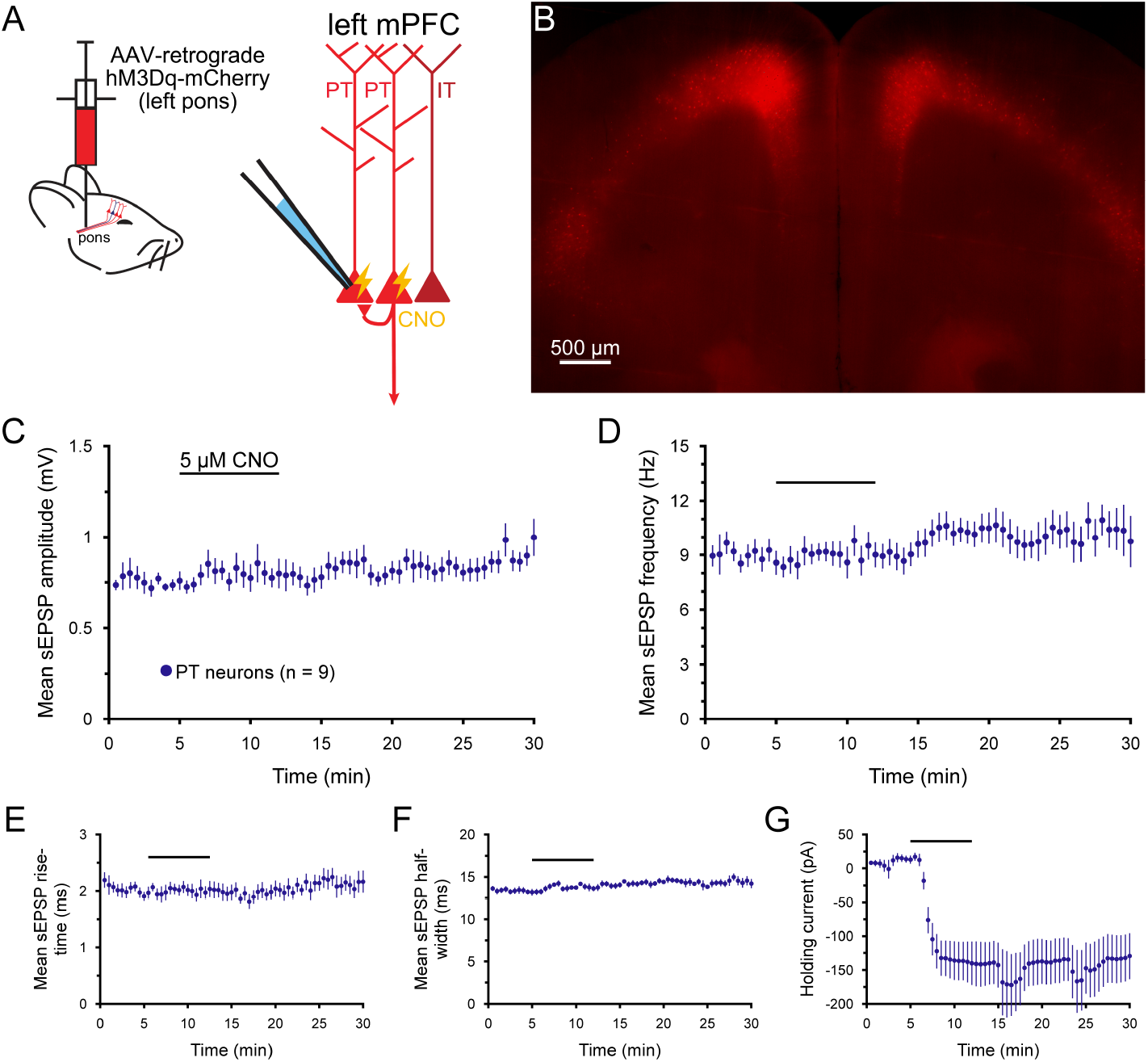
The effects of acetylcholine (ACh) on sEPSPs are not mimicked by activation of hM3Dq receptors in PT neurons. ***A***) Diagram of injection of AAV-retrograde hM3Dq-mCherry virus into the pons to express hM3Dq receptors selectively in PT neurons in the cortex (left), and of the recording setup using clozapine N-oxide (CNO; 5 μM) to excite PT neurons (right). ***B***) Image of hM3Dq-m-Cherry-expressing layer 5 PT neurons in the prefrontal cortex (5x objective, artificially colored) three weeks after virus injection into the pons. ***C***) Plot of mean sEP-SP amplitude (average of 30 second means ± SEM) over time for 9 PT neurons express-ing hM3Dq-mCherry. CNO was bath-applied for 7 minutes, as indicated. ***D*** - ***G***) Similar to ***C***, with plots of mean sEPSP frequency (***D***), rise-time (***E***), and half-width (***F***) over time. ***G***) Plot of the mean holding current necessary to maintain neurons at their initial resting membrane potentials. Note that CNO induced large and irreversible inward currents, but had no effect on sEPSP properties, including sEP-SP rise-times (***E***).

**Table 12.** Effects of chemogenetic activation of PT neurons on sEPSPs (n = 9 PT neurons)

| Measurement | Baseline | + CNO | Wash | Change in CNO | CNO vs baseline* (p-value) | Effect size (Cohen's d) |
| --- | --- | --- | --- | --- | --- | --- |
| sEPSP amplitude (mV) | 0.75 ± 0.06 | 0.80 ± 0.07 | 0.94 ± 0.7 | +5 ± 6% | 0.422 | 0.24 |
| sEPSP frequency (Hz) | 8.9 ± 0.6 | 9.3 ± 0.8 | 10.1 ± 1.3 | 6 ± 9% | 0.656 | 0.16 |
| sEPSP rise-time (ms) | 1.94 ± 0.08 | 2.01 ± 0.13 | 2.16 ± 0.19 | +3 ± 4% | 0.398 | 0.20 |
| sEPSP half-width (ms) | 13.2 ± 0.3 | 13.7 ± 0.4 | 14.4 ± 0.5 | +4 ± 2% | 0.110 | 0.45 |
| Holding current (pA) | +13 ± 4 | -139 ± 46 | -131 ± 44 | -152 ± 29 pA | < 0.001 | **2.28** |

Data are shown as means ± SEM. \*Student's t-test for paired samples. Large effect sizes ( $\geq \sim 0.80$ ) are shown in bold.

## Discussion

### Acetylcholine enhances excitatory drive preferentially in PT neurons

Given the wealth of data demonstrating cholinergic suppression of glutamate release at excitatory cortical synapses (see **Introduction**), ACh was expected to reduce spontaneous excitatory synaptic drive in both IT and PT neurons. The results described above demonstrate that this is not the case. Instead, in two independent data sets, ACh promoted excitatory drive preferentially in PT neurons while having little impact on sEPSP frequencies or amplitudes in IT neurons (see **Figures 3 and 8**). The preferential impact of ACh on sEPSPs in PT neurons was robust across sEPSP detection thresholds and did not depend on the kinetics of the EPSP template used to detect events (see **Figure 3C**,**D**). The sensitivity of cholinergic enhancement of sEPSP amplitudes and frequencies to TTX (see **Figure 7**) suggests that ACh acts at the network level to enhance excitatory drive preferentially in PT neurons.

Cholinergic facilitation of excitatory drive in PT neurons is noteworthy because ACh also preferentially enhances PT neuron excitability, as reflected in the larger holding currents necessary to keep PT neurons at their initial resting membrane potentials (e.g., **Figures 3H and 8H**), a result consistent with prior reports of selective cholinergic excitation of PT neurons (Dembrow et al., 2010; Joshi et al., 2016; Baker et al., 2018b). The present results are also consistent with those of Shirey et al. (2009), who found that M1 receptor activation enhanced the amplitude and frequency of spontaneous, but not miniature, EPSPs in layer 5 neurons in the mPFC of young (<4-week-old) rats. The present study goes further in demonstrating that the cholinergic increase in excitatory drive is preferential to PT neurons and requires recurrent excitation within networks of PT neurons (see **Figures 8, 9**, and **10**). Thus, ACh release in the cortex engages two parallel mechanisms that promote corticofugal output to the brainstem: preferential increase of PT neuron excitability, and robust recurrent excitatory drive within PT networks.

Cholinergic enhancement of synaptic drive in PT neurons is attributed to M1-type muscarinic receptor activation, as it was blocked by both atropine (a broad spectrum mAChR antagonist; see **Figure 4**) and pirenzepine (an M1-selective antagonist; see **Figure 5** and Shirey et al., 2009). It is notable that, after blockade of M1 receptors, ACh moderately *decreased* sEPSP frequency in PT neurons, a result that aligns with the well-established M4-dependent presynaptic inhibition of glutamate release at neocortical (Eggermann and Feldmeyer, 2009; Yang et al., 2020) and hippocampal (Kimura and Baughman, 1997; Shirey et al., 2008; Dasari and Gulledge, 2011) synapses. Cholinergic enhancement of PT-PT recurrent activity occurring in parallel with generalized suppression of other glutamatergic afferents is expected to further focus corticofugal output in populations of synaptically-coupled PT neurons.

Activation of mAChRs can also increase the amplitude and frequency of action-potential-dependent inhibitory postsynaptic currents (IPSCs) in cortical pyramidal neurons (Kawaguchi, 1997; Kondo and Kawaguchi, 2001), potentially via direct excitation of somatostatin-expressing GABAergic interneurons (Kawaguchi, 1997; Fanselow et al., 2008; Xu et al., 2013; Chen et al., 2015; Obermayer et al., 2018). On the other hand, ACh can suppress GABA release at perisomatic inhibitory synapses made by fast-spiking interneurons (Kruglikov and Rudy, 2008)(see also, Aramakis et al., 1997; Salgado et al., 2007; Nunez et al., 2012). These results suggest that ACh may enhance inhibition to tuft dendrites while simultaneously disinhibiting perisomatic regions of projection neurons. In the present study, cholinergic enhancement of sEPSPs was similar in control conditions with intact GABAergic networks and in conditions in which GABAergic synaptic transmission was blocked (see **Figure 6**). This suggests that cholinergic modulation of inhibitory circuits is not required to enhance the excitatory synaptic drive of PT neurons in the adult prefrontal cortex. As most of the prior studies described above utilized immature (pre-weening) rodents (but see Chen et al., 2015), it is possible that there are developmental changes in the cholinergic sensitivity of inhibitory cortical circuits. Alternatively, because PT neurons typically establish perisomatic connections to other PT neurons in the local network (Morishima et al., 2011), and inputs from ACh-sensitive somatostatin neurons typically occur further out in apical dendrites (Silberberg and Markram, 2007), there may be limited interaction among the two inputs. Regardless, enhanced inhibition of the apical tuft would be expected to further shift the weight of excitatory drive toward more proximal PT-PT recurrent connections.

Although pyramidal neurons in the mPFC lack significant postsynaptic nicotinic responses to ACh (Hedrick and Waters, 2015), presynaptic nAChRs may modulate glutamate (Lambe et al., 2003; Couey et al., 2007; Aracri et al., 2013) and GABA (Couey et al., 2007; Aracri et al., 2010) release onto layer 5 neurons in the dorsal mPFC. In addition, ACh may excite a subpopulation of cortical GABAergic interneurons that express postsynaptic nAChRs (Poorthuis et al., 2014). The present study tested the role of ACh, rather than specific cholinergic agonists, in an effort to capture the net effect of ACh on network activity in IT and PT neurons, regardless of which receptor subtypes might be involved (i.e., mAChRs or nAChRs). However, the ability of atropine to completely block all cholinergic effects on sEPSPs (**Figure 4**) suggests that nAChRs may not be effective in regulating local network excitatory drive to adult IT and PT neurons (see also Aramakis and Metherate, 1998).

### Acetylcholine promotes recurrent activity in networks of PT neurons

Simultaneous recordings in pairs of neurons allowed direct comparisons of net excitatory drive to IT and PT neurons, and for the detection and measurement of synchronous sEPSPs under different experimental conditions. In baseline conditions the rate of synchronous sEPSPs was very low, either equivalent to random chance (IT-IT pairs) or somewhat above random chance (in IT-PT and PT-PT pairs). ACh preferentially increased the rate of synchronous sEPSPs in PT-PT pairs (see **Figure 8**; **Table 8**). Given that TTX reduced sEPSP frequencies in PT neurons by ∼25% (see **Figure 7**), it can be estimated that mEPSPs account for the remaining ∼75% of events. The abundance of mEPSPs, which by nature are not purposefully synchronized across release sites, and which are not enhanced by ACh, likely occludes the full impact of cholinergic synchronization of action-potential-evoked sEPSPs in PT neurons. Therefore, it is plausible that cholinergic stimulation will synchronize upwards of ∼8% of action-potential-dependent sEPSPs in individual PT-PT pairs. The additional excitatory drive to PT neurons in the presence of ACh is attributable to recurrent PT networks, as it was eliminated by chemogenetic silencing of PT, but not IT, neurons (see **Figures 9** and **10**). Although chemogenetic excitation of PT neurons failed to mimic cholinergic enhancement of sEPSPs (**Figure 11**), it also failed to reproduce other aspects of cholinergic signaling (e.g., the increase in sEPSP rise-times), making it difficult to interpret this negative finding. Alternative approaches to selectively activate PT networks will be needed to confirm whether increased PT activity is sufficient to enhance sEPSP amplitudes and frequencies.

Despite a lack of *in vivo* data regarding synchronized excitatory input to PT neurons, it is notable that populations of motor cortex PT neurons that share projection targets exhibit concurrent activity during motor behavior (Nelson et al., 2021)(see also, Olivares-Moreno et al., 2019), raising the possibility that they also share excitatory drive. Given that PT neurons selectively form strong unitary connections with other PT neurons (Morishima and Kawaguchi, 2006; Morishima et al., 2011), and that cortical ACh release is associated with the initiation of motor activity (Day et al., 1991; Parikh et al., 2007; Jing et al., 2020; Lohani et al., 2022), ACh may act to coordinate activity in ensembles of PT neurons to guide behavior.

### Conclusions and considerations

The results of this study demonstrate that ACh, acting on M1-type mAChRs, promotes corticofugal output of the prefrontal cortex by enhancing recurrent excitation in networks of PT neurons. This study was focused on comparing the net impact of ACh on local network synaptic drive to IT an PT neurons. However, many excitatory synapses arise from long-distance corticocortical and thalamocortical afferents that are not expected to contribute to action-potential-dependent synaptic signaling in the *ex vivo* conditions utilized here. Therefore, to fully elucidate how ACh regulates information flow in the neocortex it will be important for future studies to test the impact of ACh on evoked transmitter release from a range of cortical afferents. This has been done to a limited extent in the hippocampus, where differential presynaptic sensitivity to ACh is thought to shift the weight of synaptic drive from Shaffer collateral (CA3-to-CA1) inputs toward more distal excitatory inputs from the entorhinal cortex (Hasselmo and Schnell, 1994; Thorn et al., 2017). While there are some data suggestive of similar afferent-specific cholinergic modulation in the neocortex (Gil et al., 1997; Hsieh et al., 2000; Lambe et al., 2003; Yang et al., 2020), a more complete understanding of how ACh regulates cortical output will require experiments that test for afferent-specific cholinergic modulation across a range of isolated local and long-distance excitatory inputs to IT and PT neurons.

## Acknowledgements

I thank Saiko Ikeda for technical assistance with intracranial virus injections and data analysis, and Arielle L. Baker for assistance with generating a physiology index to distinguish IT and PT neuron subpopulations.

## Notes

### Competing Interest Statement

The authors have declared no competing interest.

### Summary of Updates

1) The data for synchronous input to homotypic and heterotypic neuron pairs was reanalyzed pairwise, and Figure 8C and Table 8 updated to reflect those new analyses. 2) The text of the manuscript was updated to reflect the new analyses for Figure 8C and Table 8.

